# Modeling and Characterization of Inter-Individual Variability in CD8 T Cell Responses in Mice

**DOI:** 10.1101/671891

**Authors:** Chloe Audebert, Daphné Laubreton, Christophe Arpin, Olivier Gandrillon, Jacqueline Marvel, Fabien Crauste

**Author notes:** These authors contributed equally to this work (Jacqueline Marvel, Fabien Crauste).

## Abstract

To develop vaccines it is mandatory yet challenging to account for inter-individual variability during immune responses. Even in laboratory mice, T cell responses of single individuals exhibit a high heterogeneity that may come from genetic backgrounds, intra-specific processes (*e*.*g*. antigen-processing and presentation) and immunization protocols.

To account for inter-individual variability in CD8 T cell responses in mice, we propose a dynamical model coupled to a statistical, nonlinear mixed effects model. Average and individual dynamics during a CD8 T cell response are characterized in different immunization contexts (vaccinia virus and tumor). On one hand, we identify biological processes that generate inter-individual variability (activation rate of naive cells, the mortality rate of effector cells, and dynamics of the immunogen). On the other hand, introducing categorical covariates to analyze two different immunization regimens, we highlight the steps of the response impacted by immunogens (priming, differentiation of naive cells, expansion of effector cells and generation of memory cells). The robustness of the model is assessed by confrontation to new experimental data.

Our approach allows to investigate immune responses in various immunization contexts, when measurements are scarce or missing, and contributes to a better understanding of inter-individual variability in CD8 T cell immune responses.

## 1 Introduction

The immune response is recognized as a robust system able to counteract invasion by diverse pathogens [1, 2]. However, as a complex biological process, the dynamical behavior of its cellular components exhibits a high degree of variability affecting their differentiation, proliferation or death processes. Indeed, the abundance of antigen-specific T cells and their location relative to pathogen invasion will affect the dynamic of the response [2–4]. Similarly, the pathogen load and virulence as well as the host innate response will affect the T cell response [5]. Finally, at the cellular level, between-cell variations in protein content can also generate heterogeneous responses [6]. Genetic variability of the numerous genes controlling the immune response will also be a source of variability among individuals [1]. Even among genetically identical individuals, the response to the same infection can result in highly heterogeneous dynamics [7–9].

Cytotoxic CD8 T cells play an essential role in the fight against pathogens or tumors as they are able to recognize and eliminate infected or transformed cells. Indeed, following encounter of antigen-presenting cells loaded with pathogen or tumor derived antigens, in lymphoid organs, quiescent naive CD8 T cells will be activated. This leads to their proliferation and differentiation in effector cells that have acquired the capacity to kill their targets, and to their ultimate differentiation in memory cells [10, 11]. The CD8 T cell immune response is yet a highly variable process, as illustrated by experimental measurements of cell counts: dynamics of the responses (timing, cell counts) may differ from one individual to another [4, 12, 13], but also depending on the immunogen [3, 7, 9].

The role of genome variability in explaining inter-individual variations of T cell responses has been recently investigated [14,15] but provided limited understanding of the observed heterogeneity. Li et al. [15] focused on correlations between gene expression and cytokine production in humans, and identified a locus associated with the production of IL-6 in different pathogenic contexts (bacteria and fungi). Ferraro et al. [14] investigated inter-individual variations based on genotypic analyses of human donors (in healthy and diabetic conditions) and identified genes that correlate with regulatory T cell responses.

To our knowledge, inter-individual variability characterized by heterogeneous cell counts has been mostly ignored in immunology, put aside by focusing on average behaviors of populations of genetically similar individuals. The use of such methodology, however, assumes that variability is negligible among genetically similar individuals, which is not true [7, 10, 16], see Figure 1. Experimental measurements of *in vivo* immune responses are often limited, due to tissue accessibility. For instance, mice have to be killed to count CD8 T cell numbers in organs, restricting the follow-up of one individual to blood sampling. Also, ethics do not allow everyday bleeding of animals. Hence measurements in peripheral blood are often performed on a restricted number of time points per individual, which probably led to focus more on average dynamics rather than on heterogeneous individual dynamics.

**Figure 1.**
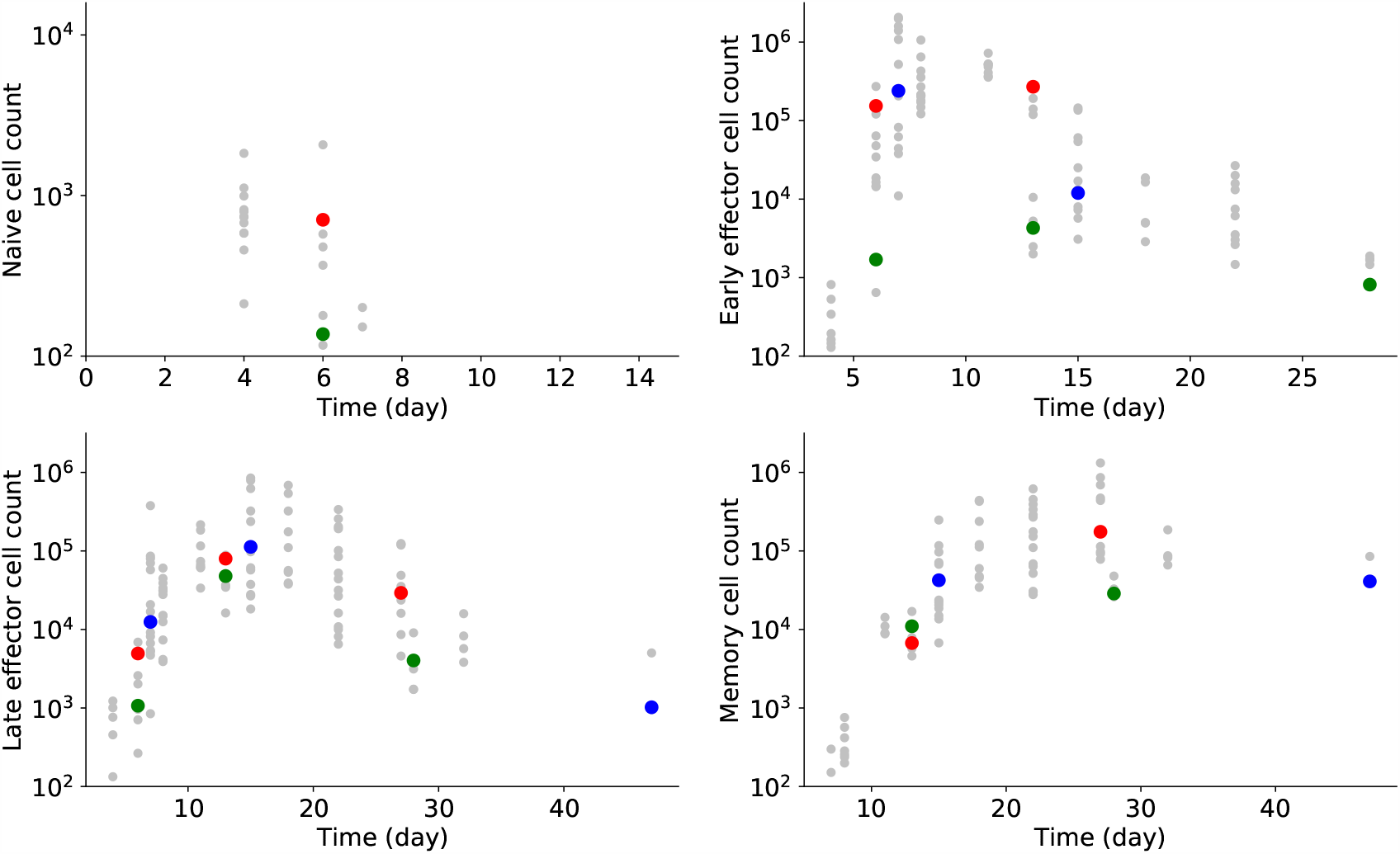
CD8 T cells counts after vaccinia virus (VV) immunization in mice. Naive CD44-Mki67-Bcl2+ cell counts (upper left), Early Effector CD44+Mki67+Bcl2-cell counts (upper right), Late Effector CD44+Mki67-Bcl2-cell counts (lower left), and Memory CD44+Mki67-Bcl2+ cell counts (lower right) are measured in 59 individuals over 47 days. Measurements of three different mice are highlighted in blue, green and red, whenever they are available. All other measurements are in grey.

In this work, we propose to model and characterize inter-individual variability based on CD8 T cell counts using nonlinear mixed effects models [17–19]. In these models, instead of considering a unique set of parameter values as characteristic of the studied data set, a so-called *population approach* is used based on distributions of parameter values. All individuals are assumed to be part of the same population, and as so they share a common behavior (average behavior) while they keep individual behaviors (random effects). Nonlinear mixed effects models are well adapted to repeated longitudinal data. They aim at characterizing and understanding “typical behaviors” as well as inter-individual variations. T cell count measurements, obtained over the course of a response (few weeks), and the large variability they exhibit represent a case study for this approach (see Figure 1).

Nonlinear mixed effects models have been used to analyze data in various fields [20], especially in pharmakokinetic studies, and more recently to model cell to cell variability [21, 22] or to study tumor growth [23, 24]. In immunology, Keersmaekers et al. [25] have recently studied the differences between two vaccines with nonlinear mixed effects models and ordinary differential equation (ODE) models for T and B cells. Jarne et al. [26] and Villain et al. [27] have used the same approach to investigate the effect of IL7 injections on HIV+ patients to stimulate the CD4 T cell response. None of these works aimed at identifying immunological heterogeneous processes or characterizing the between-individual variability in CD8 T cell responses, rather nonlinear mixed effects models have been used to characterize the average behavior of the cell populations and explain the data.

In order to characterize inter-individual variability based on experimental data of cell counts, other methods could be considered. First, one could try to estimate individual parameter values by fitting a mathematical model to individual experimental data. The nature of the data we consider here, illustrated in Figure 1, makes this option unrealistic (not enough data per individual). Another approach could be to use individual-based models, yet such models also require to estimate a lot of parameters and individual data in our case do not provide enough information *per se*. Consequently, nonlinear mixed effects models appear to be the most appropriate method to handle sparse individual data.

A number of models of the CD8 T cell response based on ODEs have been proposed over the years. Of particular relevance here is the work of De Boer et al. [28], where the model accounts for activated and memory cell dynamics but the influence of the immunogen is imposed. Antia et al. [29] proposed a model based on partial differential equations, that includes immunogen effects and dynamics of naive, effector and memory cells. These works describe different subpopulations of CD8 T cells, however most of the time only total CD8 T cell counts are available to validate the models. In Crauste et al. [10], the authors generated cell counts for four subpopulations of CD8 T cells in mice that they used to identify the most likely differentiation pathway of CD8 T cells after immunogen presentation. This approach has led to a model of the average CD8 T cell dynamics in mice after immunization and its representation as a set of nonlinear ODEs. The model consists in a system of ODEs describing the dynamics of naive, early effector, late effector, and memory CD8 T cell subsets and the immunogen.

The goal of this article is to propose, analyze and validate a mathematical methodology for describing individual behaviors and the inter-individual variability observed in CD8 T cell responses, in different immunization contexts. We will rely on a dynamical description of CD8 T cell dynamics, based on a nonlinear model, where parameter values are drawn from probability distributions (nonlinear mixed effects model). Starting from the model introduced and validated in [10], we first select a model of the CD8 T cell immune response dynamics accounting for variability in cell counts by using synthetic then experimental data, generated in different immunization contexts. Second we characterize the main biological processes contributing to heterogeneous individual CD8 T cell responses. Third, we establish that the immunogen-dependent heterogeneity influences the early phase of the response (priming, activation of naive cells, cellular expansion). Finally, we show that besides its ability to reproduce CD8 T cell response dynamics our model is able to predict individual dynamics of responses to similar immunizations, hence providing an efficient tool for investigating CD8 T cell dynamics and inter-individual variability.

## 2 Results

CD8 T cell counts have been previously experimentally measured and characterized in 4 sub-populations in Crauste et al. [10]: naive (N), early effector (E), late effector (L) and memory (M) cells (see Section 4.2). Additionally, a mathematical model has been proposed for the description of these data and validated in [10]. Therein, System (3) has been shown to be able to describe average dynamics of CD8 T cell immune responses, when CD8 T cells go through the 4 above-mentioned differentiation stages. In order to describe individual CD8 T cell counts we couple System (3) to a nonlinear mixed effects model that accounts for both population and individual dynamics. This results in a model of CD8 T cell response dynamics that includes more parameters, consequently when fitting this model to new data sets it is mandatory to determine whether all model parameters can be correctly estimated or if some parameters have to be removed and the initial model modified. We first perform this analysis on ideal data, called “synthetic data”, to determine the minimal number of parameters required to fit data with the model. Synthetic data are generated from simulations of the model, so true parameter values are known and the estimation procedure is performed in a controlled framework. Then, in a second time, we perform again the analysis on real, experimental data, in order to adapt our procedure to realistic data. Finally, we characterize the biological processes depending on the immunization, and we identify model parameters and their corresponding biological processes that vary the most between individuals. Details of the methodology are presented in Sections 4.3 to 4.7.

### 2.1 Model selection on synthetic data

In [10], System (3) introduced in Section 4.3 has been shown to be able to describe average dynamics of CD8 T cell immune responses, when CD8 T cells go through 4 differentiation stages: naive, early-then late-effector cells, and memory cells (see Section 4.2). Here System (3) is coupled to a non-linear mixed effect model, leading model parameters to be drawn from a probability distribution. Initially we assume that any parameter can carry inter-individual variability, then the number of parameters is reduced to ensure correct estimations on ideal data. Ideal data are generated by simulating System (3), so parameter values are known (probability distributions). These data sets enable to evaluate the potential of the model without data availability-related limitations: we call them”synthetic data”. Here, synthetic data account for 100 individuals (mice) and measurements are available every day from D4 to D10, then every 2 days from D10 to D20, and finally on D25 and D30 pi, see Table 1 and Section 4.6 for details.

**Table 1.** Data sets (details in Sections 4.2, 4.3 and 4.6).

| Short Name | Description |
| --- | --- |
| VV data set 1 | CD8 T cell counts of 59 individual mice inoculated intra-nasally with $2 \times 10^5$ pfu of a vaccinia virus (VV) expressing the NP68 epitope ; naive, early and late effector, and memory cell counts have been measured up to day 47pi (days 4, 6, 7, 8, 11, 13, 15, 18, 22, 27, 28, 32, 47pi with maximum 15 measurements per time point) |
| VV data set 2 | Similar to VV data set 1 (15 individual mice) ; CD8 T cell counts of naive, early and late effector, and memory cells have been measured following VV immunization, up to day 42pi (days 4, 6, 7, 8, 11, 13, 15, 21, 28, 42pi with maximum 4 measurements per time point) |
| Tumor data set 1 | CD8 T cell counts of 55 individual mice subcutaneously inoculated with $2.5 \times 10^6$ EL4 lymphoma cells expressing the NP68 epitope ; naive, early and late effector, and memory cell counts have been measured up to day 47pi (days 4, 5, 6, 7, 8, 10, 11, 13, 14, 15, 18, 22, 27, 32, 46, 47pi with maximum 15 measurements per time point) |
| Tumor data set 2 | Similar to Tumor data set 1 (20 individual mice); CD8 T cell counts of naive, early and late effector, and memory cells have been measured following Tumor immunization, up to day 42pi (days 6, 7, 8, 11, 13, 15, 21, 28, 42pi with maximum 5 measurements per time point) |
| Synth data sets 1 to 4 | Synthetic data sets generated with System (3) and its subsequent simplifications (see Section 4.6), consisting in CD8 T cell counts of naive, early and late effector, and memory cells on days 4, 5, 6, 7, 8, 9, 10, 12, 14, 16, 18, 20, 25, 30pi for all 100 individuals |

We use Synth data sets 1 to 4 (Table 1) to validate the parameter estimation procedure. True parameter values are known and given in Section 5.1. Parameter estimation is performed with the SAEM algorithm implemented in Monolix software [30]. Using a model selection procedure, based in particular on the use of the relative standard error (RSE) defined in (4) that informs on the confidence in the estimation, we iteratively remove parameters:

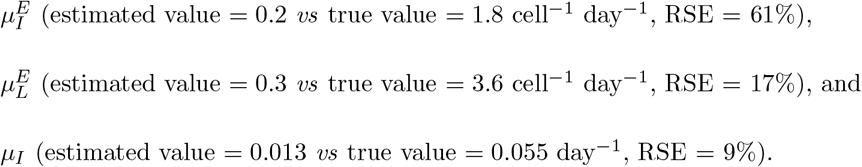

Details of the procedure are explained in Sections 4.5, 4.6, 5.2 and Table A.2.

All removed parameters are related to death rates, of late effector cells 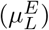 and of the immunogen 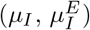). In each case, the model still accounts for death of late effector cells and of the immunogen, *via* parameters 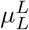 and 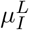. Nonlinear mixed effects models avoid redundancy in the description of biological processes, thus they allow reliable parameter estimation using synthetic data.

This leads to a reduction of the initial 12-parameters System (3) to the 9-parameters System (1),

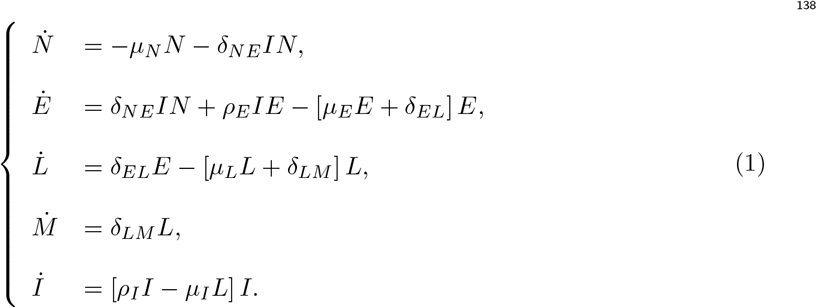

where all parameters still account for individual variability (drawn from probability distributions). For the sake of simplicity the parameters are renamed in System (1):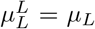 and 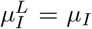. Figure 2.A displays a schematic representation of System (1).

**Figure 2.**
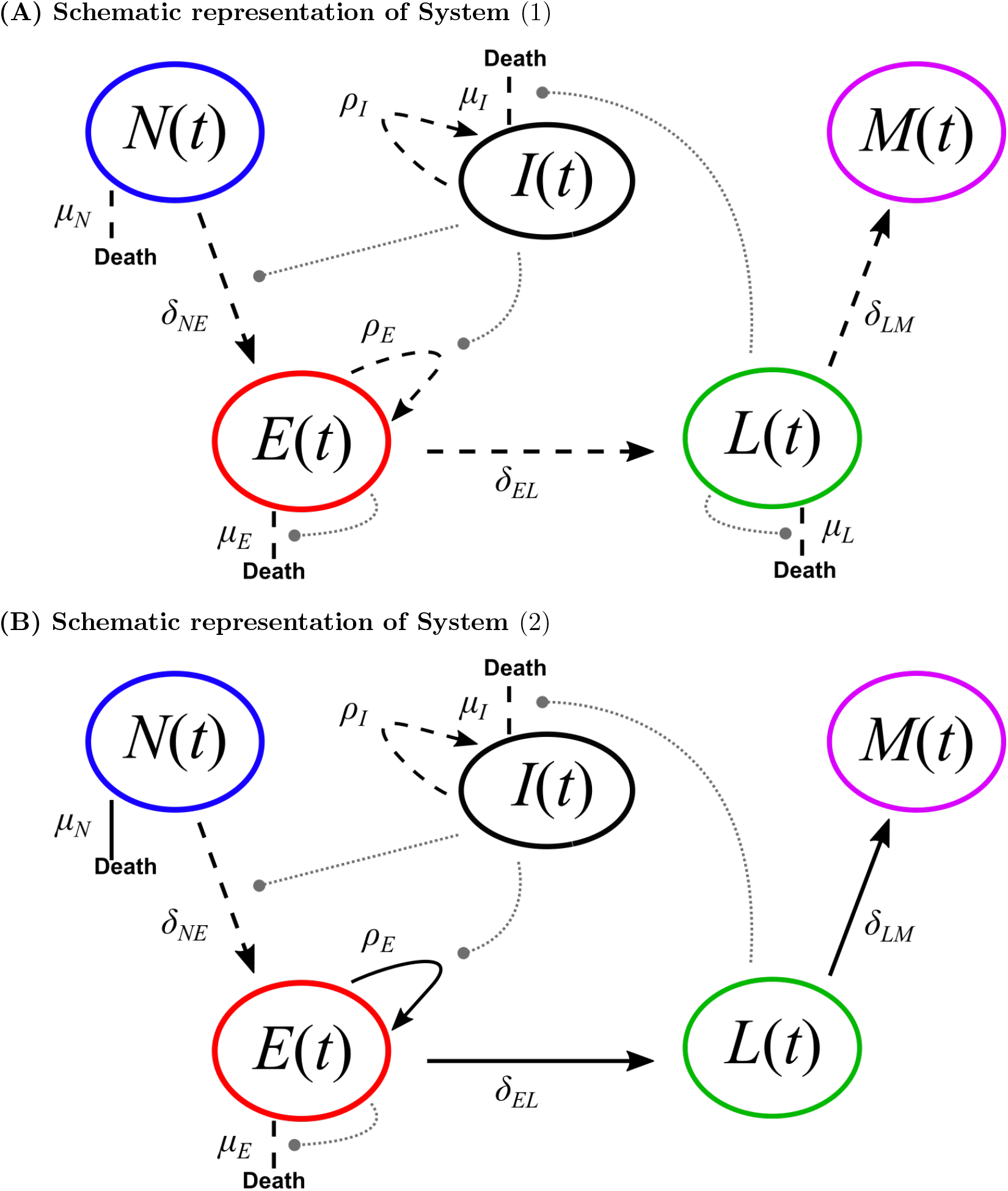
Schematic CD8 T cell differentiation diagram following immunization. **(A)** Schematic representation of System (1). **(B)** Schematic representation of System (2). Dashed black lines represent individual-dependent parameters, while straight black lines (only in **(B)**) represent parameters fixed within the population. Grey round-ended dotted lines represent feedback functions (see systems of equations).

### 2.2 A model of CD8 T cell dynamics accounting for *in vivo* inter-individual heterogeneity

Biological data from VV data set 1 (see Section 4.2 and Table 1) are confronted to System (1). Parameter estimation is performed using the SAEM algorithm [30] and, following the procedure described in Section 4.7, leads to further reduction of the model. Using *in vivo* data to estimate parameter values provides a priori less information than synthetic data: there are 59 individuals instead of 100 for synthetic data, and at most 15 measurements (15 individuals) at each time point are available whereas almost all measurements are available for the 100 individuals in synthetic data. Hence, it might be necessary to simplify the model to ensure correct parameter estimations, either mean values or random effects, similarly to what has been done in the previous section.

The first step in the model reduction procedure leads to an estimated value of parameter *µ*_*L*_ close to zero (2 × 10^*−*8^ cell^*−*1^ day^*−*1^), with a RSE *>* 100%, see Table 2, *Step 1*. Hence parameter *µ*_*L*_ is removed, and the estimation is performed again with the updated model. We observe that all mean value parameters have now RSE *<* 30%, so we conclude that their estimations are reliable (Table 2, *Step 2*).

**Table 2.** Steps in estimating parameter values using VV data set 1 and System (1). The procedure is detailed in Section 4.7. At *Step 1*, the procedure leads to removing parameter *µ*_*L*_. At *Step 2*, the random effect of *δ*_*LM*_ is removed. At *Step 3*, the random effect of *δ*_*EL*_ is removed. At *Step 4*, the random effect of *ρ*_*E*_ is removed. At *Step 5*, the random effect of *µ*_*N*_ is removed. At *Step 6*, no other action is required. Values used to take a decision are highlighted in bold at each step. In the first column, ‘m.v.’ stands for mean value, RSE is defined in (4), ‘r.e.’ stands for random effect, and the shrinkage is defined in (5). Note that values (mean values and random effects) of parameters *µ*_*E*_, *µ*_*L*_, and *µ*_*I*_ have to be multiplied by 10^*−*6^ (for *µ*_*E*_ and *µ*_*L*_) and 10^*−*5^ (for *µ*_*I*_). Units are omitted for the sake of clarity.

| | $\mu_N$ | $\delta_{NE}$ | $\rho_E$ | $\mu_E$ | $\delta_{EL}$ | $\mu_L$ | $\delta_{LM}$ | $\rho_I$ | $\mu_I$ |
| --- | --- | --- | --- | --- | --- | --- | --- | --- | --- |
| *Step 1* |  |  |  |  |  |  |  |  |  |
| m.v. | 0.59 | 0.002 | 0.9 | 5.2 | 0.13 | **0.02** | 0.09 | 0.08 | 1.9 |
| RSE | 5 | 30 | 2 | 21 | 11 | **207** | 8 | 7 | 28 |
| r.e. | 0.16 | 0.8 | 0.04 | 0.67 | 0.1 | 1.9 | 0.05 | 0.2 | 1.3 |
| RSE | 66 | 36 | 44 | 35 | 567 | 220 | $10^3$ | 25 | 18 |
| shrinkage | 82 | 76 | 97 | 77 | 98 | 99 | 100 | 62 | 45 |
| *Step 2* |  |  |  |  |  |  |  |  |  |
| m.v. | 0.60 | 0.003 | 0.9 | 4.8 | 0.12 | - | 0.10 | 0.09 | 2.3 |
| RSE | 5 | 29 | 3 | 21 | 10 | - | 8 | 6 | 25 |
| r.e. | 0.15 | 0.8 | 0.06 | 0.73 | 0.2 | - | 0.05 | 0.2 | 1.2 |
| RSE | 69 | 34 | 71 | 29 | 150 | - | **$10^3$** | 25 | 17 |
| shrinkage | 83 | 78 | 94 | 74 | 96 | - | **99** | 64 | 47 |
| *Step 3* |  |  |  |  |  |  |  |  |  |
| m.v. | 0.60 | 0.001 | 1.00 | 5.0 | 0.12 | - | 0.10 | 0.08 | 2.0 |
| RSE | 5 | 32 | 1 | 20 | 10 | - | 8 | 7 | 29 |
| r.e. | 0.16 | 0.8 | 0.04 | 0.67 | 0.2 | - | - | 0.2 | 1.3 |
| RSE | 55 | 35 | 20 | 31 | **138** | - | - | 25 | 18 |
| shrinkage | 81 | 75 | 98 | 77 | **95** | - | - | 63 | 45 |
| *Step 4* |  |  |  |  |  |  |  |  |  |
| m.v. | 0.59 | 0.002 | 0.9 | 5.2 | 0.12 | - | 0.10 | 0.09 | 2.1 |
| RSE | 5 | 29 | 2 | 20 | 11 | - | 8 | 6 | 24 |
| r.e. | 0.12 | 0.9 | 0.04 | 0.74 | - | - | - | 0.2 | 1.2 |
| RSE | 102 | 32 | **47** | 29 | - | - | - | 24 | 16 |
| shrinkage | 89 | 75 | **97** | 72 | - | - | - | 63 | 49 |
| *Step 5* |  |  |  |  |  |  |  |  |  |
| m.v. | 0.59 | 0.004 | 0.8 | 4.5 | 0.12 | - | 0.10 | 0.09 | 2.7 |
| RSE | 5 | 32 | 4 | 21 | 11 | - | 8 | 6 | 21 |
| r.e. | 0.15 | 0.8 | - | 0.85 | - | - | - | 0.2 | 1.0 |
| RSE | **72** | 32 | - | 24 | - | - | - | 23 | 16 |
| shrinkage | **84** | 73 | - | 65 | - | - | - | 61 | 57 |
| *Step 6* |  |  |  |  |  |  |  |  |  |
| m.v. | 0.60 | 0.001 | 1.0 | 5.3 | 0.12 | - | 0.10 | 0.08 | 1.9 |
| RSE | 5 | 29 | 0 | 20 | 11 | - | 8 | 7 | 27 |
| r.e. | - | 0.9 | - | 0.69 | - | - | - | 0.2 | 1.3 |
| RSE | - | 29 | - | 28 | - | - | - | 23 | 17 |
| shrinkage | - | 72 | - | 75 | - | - | - | 62 | 46 |

In the second step of the procedure however, several random effects have large RSE and high shrinkages (Table 2, *Step 2* to *Step 5*). The shrinkage is defined in (5) as a measure of the difference between the estimated variance of a parameter and the empirical variance of its random effect. Parameter *δ*_*LM*_ has the worst RSE and the largest shrinkage (99%), so we remove the random effect of *δ*_*LM*_. Estimating parameter values with the updated reduced model leads to removing successively random effects of *δ*_*EL*_ (RSE = 138%, shrinkage = 95%), *ρ*_*E*_ (shrinkage = 97%), and *µ*_*N*_ (shrinkage = 84%). At each step, RSE of mean value parameters are low, and quality of individual fits is preserved.

The resulting model, System (2) (see Figure 2.B), comprises 8 parameters, 4 of them vary within the population (*δ*_*NE*_, *µ*_*E*_, *ρ*_*I*_, *µ*_*I*_) and 4 are fixed within the population (*µ*_*N*_, *ρ*_*E*_, *δ*_*EL*_, *δ*_*LM*_):

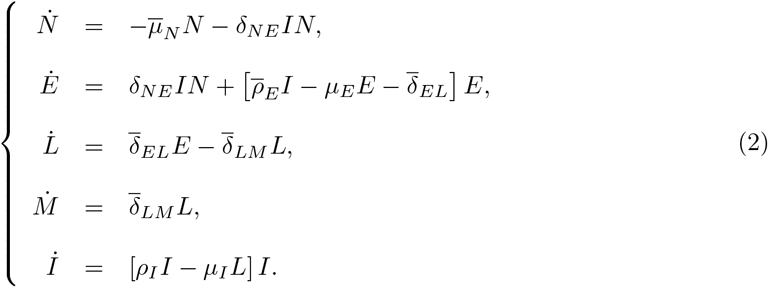

Bars highlight fixed parameters within the population. This system enables to describe VV data set 1 and its inter-individual variability (see Figure 3). The inter-individual variability is entirely contained in the activation rate of naive cells (*δ*_*NE*_), the mortality-induced regulation of effector cells (*µ*_*E*_), and the dynamics of the immunogen (*ρ*_*I*_ and *µ*_*I*_).

**Figure 3.**
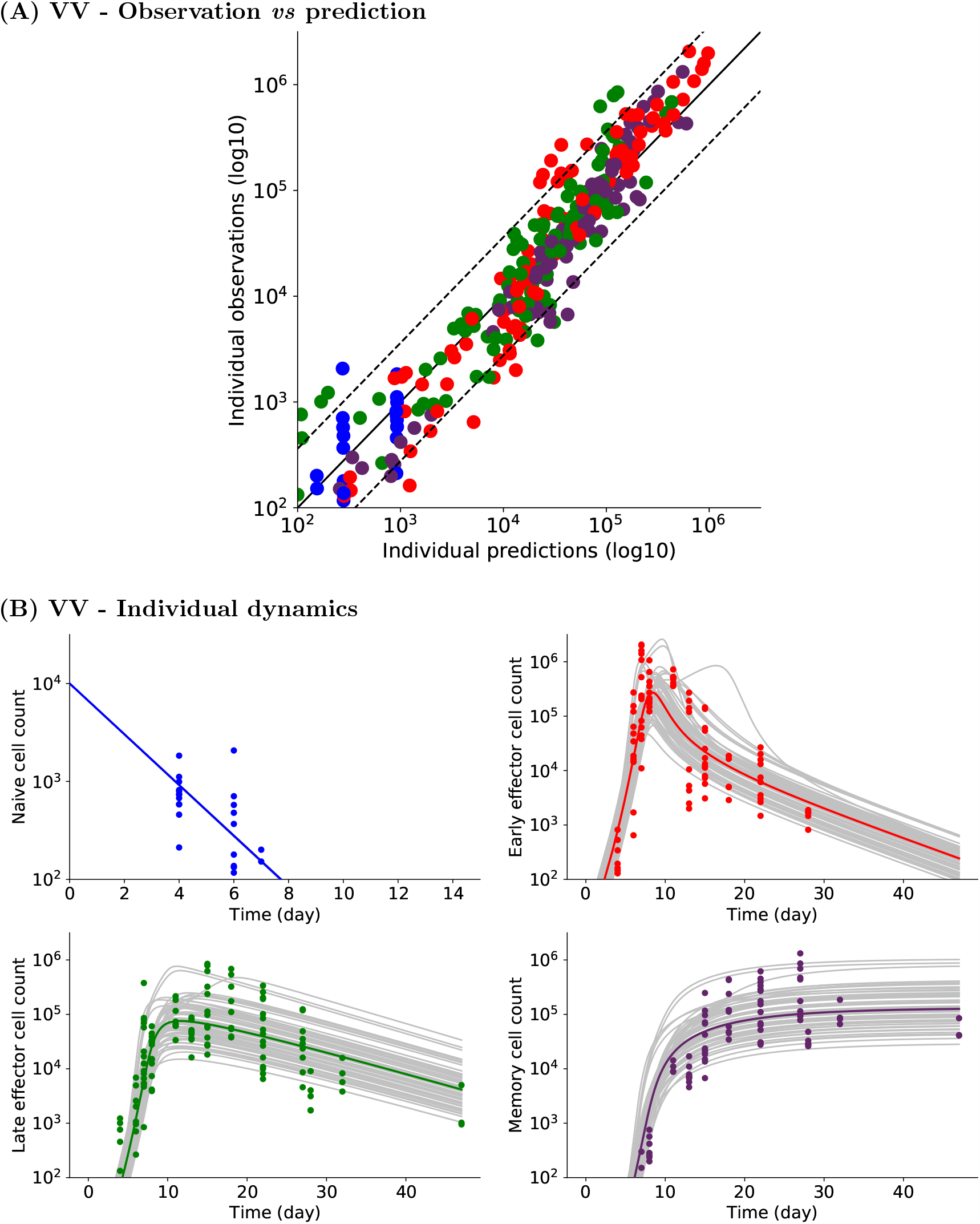
Experimental and simulated individual cell counts for VV data set 1 (logarithmic scale). **(A)** Observed *vs* predicted values. For each CD8 T cell count experimental point, the prediction is obtained by simulating System (2). Naive (blue), early effector (red), late effector (green), and memory (purple) cell counts are depicted. Dashed lines represent the 90th percentile of the difference between observed and predicted values, and the solid black line is the curve *y* = *x*. **(B)** Naive (upper left, blue), early effector (upper right, red), late effector (lower left, green) and memory (lower right, purple) cell counts up to D47pi. Experimental measurements are represented by colored dots (same color code), simulated individual trajectories by grey lines, and the average population dynamics by a straight colored line (same color code).

Figure 3.A shows the good agreement between model predictions and individual measurements for each CD8 T cell subpopulation. Model predictions are obtained from numerical simulations of System (2) performed with estimated individual parameter values. Despite over- or under-estimation of some individual observations, the 90th percentile of the difference between observed and predicted values (dashed line) shows that most experimental cell counts are efficiently predicted (estimated errors are relatively small for all subpopulations: *a*_*N*_ = *a*_*M*_ = 0.3 log10(cells), *a*_*E*_ = *a*_*L*_ = 0.4 log10(cells)). Parameter values are given in Table 3.

**Table 3.** Estimated parameter values for VV and Tumor data sets 1 (median of log-normal distribution for parameters with random effects, RSE (%) in parentheses), obtained with System (2), and estimated parameter values from [10] (VV immunization). Estimations have been performed independently.

| Parameters | Units | Estimated Values (RSE%) |  |  |  | Values from<br>Crauste et al.,<br>(2017) |
| --- | --- | --- | --- | --- | --- | --- |
|  |  | VV<br>data set 1 |  | Tumor<br>data set 1 |  |  |
| *Parameters fixed within the population* |  |  |  |  |  |  |
| $\bar{\mu}_N$ | day<sup>-1</sup> | 0.60 | (5) | 0.32 | (15) | 0.75 |
| $\bar{\rho}_E$ | day<sup>-1</sup> | 1.02 | (0) | 0.43 | (4) | 0.64 |
| $\bar{\delta}_{EL}$ | day<sup>-1</sup> | 0.12 | (9) | 0.10 | (3) | 0.59 |
| $\bar{\delta}_{LM}$ | day<sup>-1</sup> | 0.10 | (8) | 0.07 | (14) | 0.03 |
| *Parameters varying within the population* |  |  |  |  |  |  |
| $\delta_{NE}$ | day<sup>-1</sup> | 0.001 | (29) | 0.063 | (22) | 0.009 |
| $\omega_{\delta_{NE}}$ | day<sup>-1</sup> | 0.9 | (29) | 0.4 | (54) | - |
| $\mu_E$ | 10<sup>-6</sup> cell<sup>-1</sup> day<sup>-1</sup> | 5.3 | (20) | 4.9 | (18) | 21.5 |
| $\omega_{\mu_E}$ | 10<sup>-6</sup> cell<sup>-1</sup> day<sup>-1</sup> | 0.7 | (28) | 0.2 | (78) | - |
| $\rho_I$ | day<sup>-1</sup> | 0.08 | (6) | 0.11 | (3) | 0.64 |
| $\omega_{\rho_I}$ | day<sup>-1</sup> | 0.23 | (23) | 0.06 | (58) | - |
| $\mu_I$ | 10<sup>-5</sup> cell<sup>-1</sup> day<sup>-1</sup> | 1.9 | (26) | 2.4 | (18) | 1.8 |
| $\omega_{\mu_I}$ | 10<sup>-5</sup> cell<sup>-1</sup> day<sup>-1</sup> | 1.3 | (17) | 0.6 | (22) | - |
| *Residual errors* |  |  |  |  |  |  |
| $a_N$ | cell counts (log10) | 0.3 | (15) | 0.5 | (14) | - |
| $a_E$ | cell counts (log10) | 0.4 | (10) | 0.5 | (9) | - |
| $a_L$ | cell counts (log10) | 0.4 | (9) | 0.6 | (8) | - |
| $a_M$ | cell counts (log10) | 0.3 | (10) | 0.5 | (10) | - |

Figure 4 shows the estimated dynamics of early-and late-effector and memory cells of two individuals. One individual (Figure 4.A) was monitored on days 7, 15 and 47pi leading to three measurements points for late effector cells and two for early effector and memory cells. Despite missing measurements (memory cell counts on D7pi, and early effector cell counts on D47pi), estimated dynamics are in agreement with what is expected, especially regarding the time and height of the peak of the response and the following contraction phase. The other individual (Figure 4.B) had cell count measurements only on day 8pi, yet the estimated dynamics correspond to an expected behavior. This could not have been obtained by fitting this individual alone. Hence we are able to simulate likely dynamics even with a small amount of data points and missing cell count measurements at some time points, thanks to the use of nonlinear mixed effects models and the parameter estimation procedure. By focusing first on the population dynamics (based on a collection of individual dynamics), the method enables to recover the whole individual dynamics. This is a huge advantage when data sampling frequency is low.

**Figure 4.**
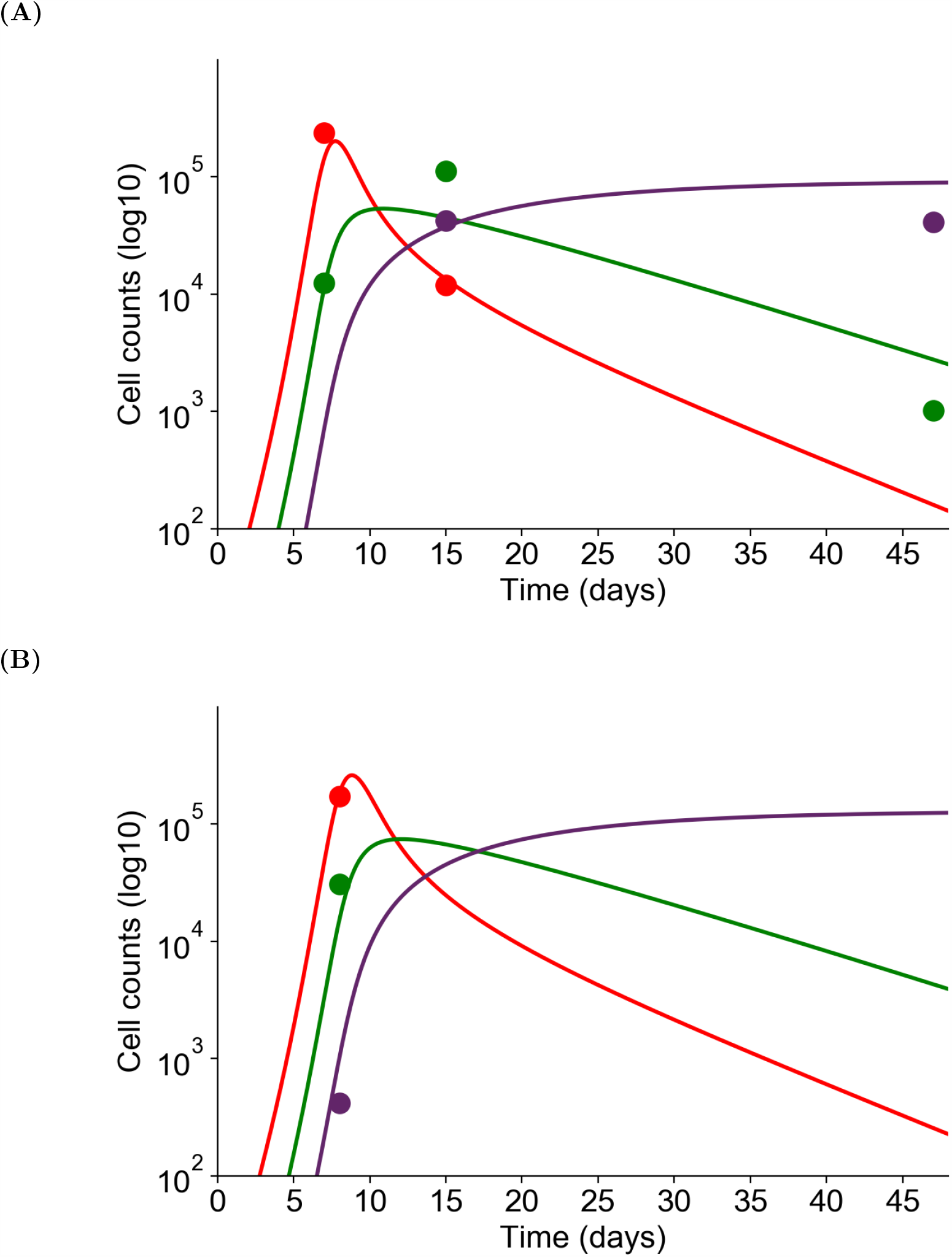
The dynamics of three subpopulations (early effector - red, late effector - green, memory - purple) are simulated with System (2) for two individuals. Experimental measurements are represented by dots, simulations of the model by straight lines. **(A)** Individual cell counts have been measured on days 7, 15 and 47pi. **(B)** Individual cell counts have been measured on day 8pi only. Although each individual is not characterized by enough experimental measurements to allow parameter estimation on single individuals, nonlinear mixed effects models provide individual fits by considering a population approach.

Similar good results are obtained for Tumor data set 1 (see Figure 5 and parameter values in Table 3). Therefore System (2) enables to describe inter-individual variability in different immunization contexts, here VV and Tumor immunizations, and with only few data points per individual.

**Figure 5.**
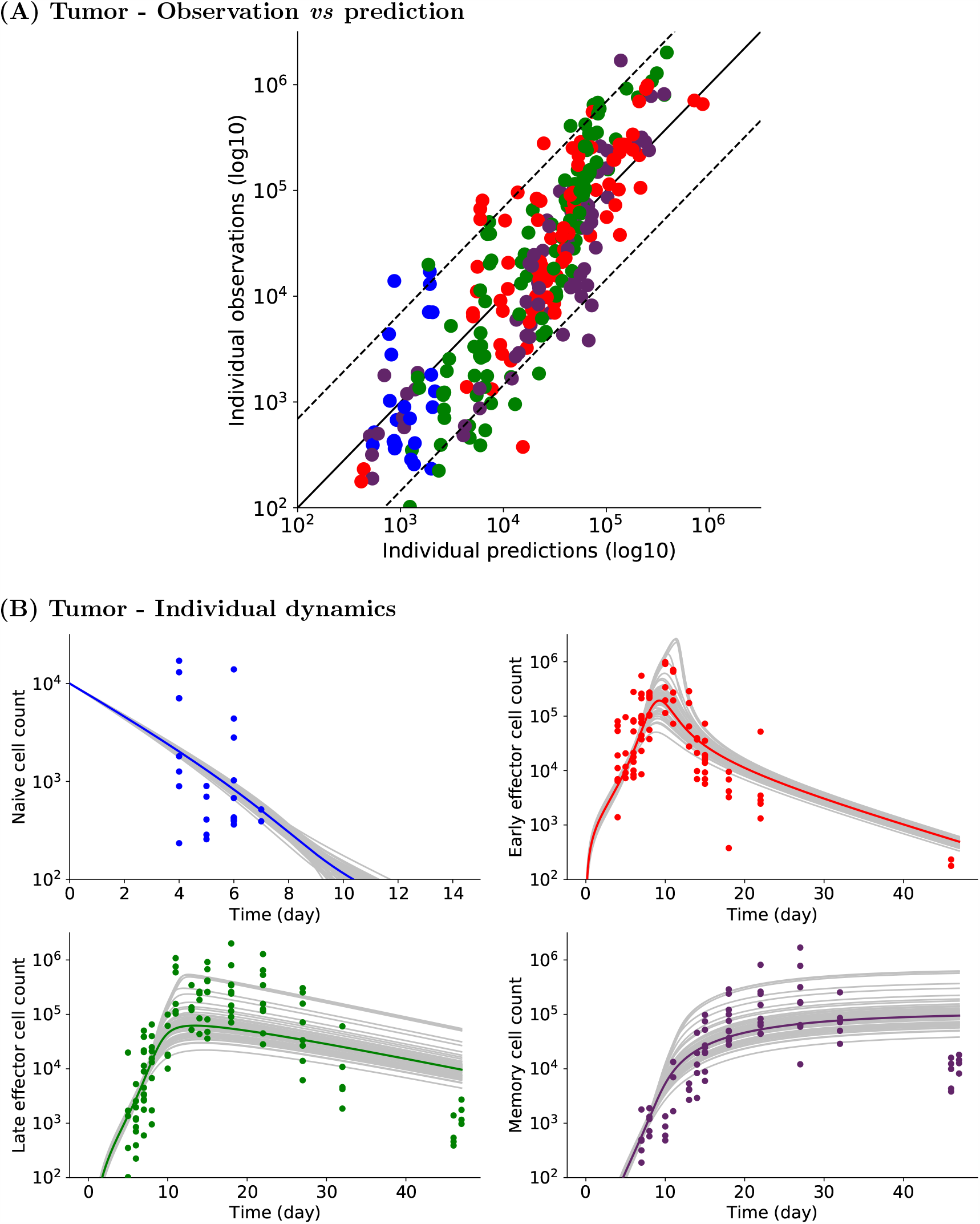
Experimental and simulated individual cell counts for Tumor data set 1 (logarithmic scale). **(A)** Observed *vs* predicted values. For each CD8 T cell count experimental point, the prediction is obtained by simulating System (2). Naive (blue), early effector (red), late effector (green), and memory (purple) cell counts are depicted. Dashed lines represent the 90th percentile of the difference between observed and predicted values, and the solid black line is the curve *y* = *x*. **(B)** Naive (upper left, blue), early effector (upper right, red), late effector (lower left, green) and memory (lower right, purple) cell counts up to D47pi. Experimental measurements are represented by colored dots (same color code), simulated individual trajectories by grey lines, and the average population dynamics by a straight colored line (same color code).

Estimated parameter values obtained with System (2) for VV or Tumor data sets are in the same range as in the estimation previously performed on average cell counts on a similar experimental set (VV immunization [10]), see Table 3. Some differences are observed for estimated values of differentiation rates, yet for the 3 estimations (VV data set 1, Tumor data set 1, [10]) parameter values remain in the same order of magnitude, indicating consistency between the two studies. Estimated values of parameter 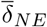 show the largest relative differences. Yet, the largest difference is observed between VV and Tumor data sets obtained with System (2), rather than between these values and the one obtained in [10]. This may highlight a potential difference in the capacity of the two immunogens (VV and Tumor) to activate naive cells. This is investigated in the next section.

### 2.3 Immunization-dependent parameters

#### Parameter comparison between immunizations

VV and Tumor induced immunizations differ in many aspects. VV immunizations are virus-mediated, use the respiratory tract to infect cells, and trigger an important innate response. Tumor immunizations involve eukaryotic cells bearing the same antigen, use subcutaneous routes, and induce a reduced innate response.

From the independent estimations on VV and Tumor data sets (Table 3), we compute differences between estimated values of fixed effects. Differences are large for parameters – in decreasing order – *δ*_*NE*_ (62%), 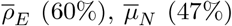, *ρ*_*I*_ (37%), and 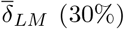. These large differences may result from biological processes involved in the CD8 T cell response that differ depending on the immunogen.

Consequently, combining both data sets (VV and Tumor) as observations may highlight which parameters have to be significantly different to describe both data sets.

#### Parameters depending on immunization

To perform this analysis, we combine VV and Tumor data sets 1 and we include a categorical covariate into the model to estimate parameter values (see Section 4.5). Covariates allow to identify parameter values that are significantly different between two CD8 activation conditions (tumors *vs* virus).

A covariate is added to the fixed effects of the five parameters that showed the larger differences in the initial estimation: *δ*_*NE*_, 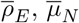, *ρ*_*I*_ and 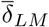. This results in the estimation of two different parameter values for parameters 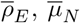 and 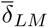 (that are fixed within the population) and two probability distributions with different mean values for parameters *δ*_*NE*_ and *ρ*_*I*_ (that vary within the population).

One may note that adding a covariate increases the number of parameters to estimate. However, the number of parameters is not doubled, since we assumed that parameters without covariates are shared by both immunization groups. In addition, the data set is larger, since it combines VV and Tumor measurements. Hence the number of parameters with respect to the amount of data remains reasonable.

From this new estimation, we conclude that among the five selected parameters the covariates of only four of them are significantly different from zero: *δ*_*NE*_, 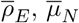, and 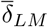 (Wald test, see Section 4.5). The estimation is therefore performed a second time assuming *ρ*_*I*_ distribution is the same in both groups. Then the Wald test indicates that the remaining covariates are significantly different from zero (Table 4).

**Table 4.**
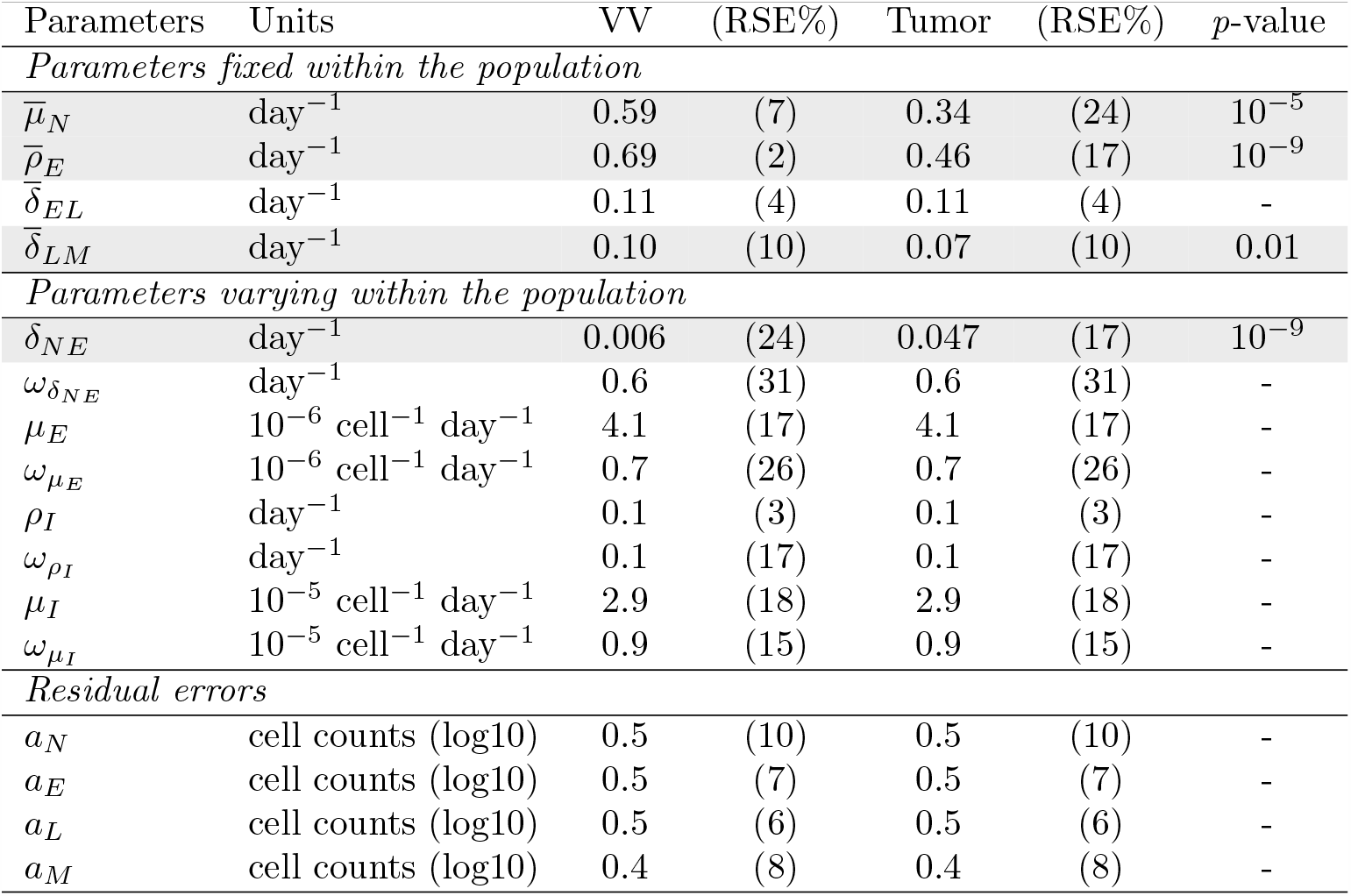
Estimated parameter values using combined VV and Tumor data sets 1. Parameters that do not vary within the population are shown in the upper part of the table, whereas individual-dependent parameters are shown in the central part (mean and standard deviation values). RSE (%) are indicated in parentheses. Parameters whose values depend on the immunogen (VV, Tumor) are highlighted in grey, and the *p*-value characterizing the covariate non-zero value is shown in the last column.

| Parameters | Units | VV | (RSE%) | Tumor | (RSE%) | $p$-value |
| --- | --- | --- | --- | --- | --- | --- |
| *Parameters fixed within the population* |  |  |  |  |  |  |
| $\bar{\mu}_N$ | day<sup>-1</sup> | 0.59 | (7) | 0.34 | (24) | 10<sup>-5</sup> |
| $\bar{\rho}_E$ | day<sup>-1</sup> | 0.69 | (2) | 0.46 | (17) | 10<sup>-9</sup> |
| $\bar{\delta}_{EL}$ | day<sup>-1</sup> | 0.11 | (4) | 0.11 | (4) | - |
| $\bar{\delta}_{LM}$ | day<sup>-1</sup> | 0.10 | (10) | 0.07 | (10) | 0.01 |
| *Parameters varying within the population* |  |  |  |  |  |  |
| $\delta_{NE}$ | day<sup>-1</sup> | 0.006 | (24) | 0.047 | (17) | 10<sup>-9</sup> |
| $\omega_{\delta_{NE}}$ | day<sup>-1</sup> | 0.6 | (31) | 0.6 | (31) | - |
| $\mu_E$ | 10<sup>-6</sup> cell<sup>-1</sup> day<sup>-1</sup> | 4.1 | (17) | 4.1 | (17) | - |
| $\omega_{\mu_E}$ | 10<sup>-6</sup> cell<sup>-1</sup> day<sup>-1</sup> | 0.7 | (26) | 0.7 | (26) | - |
| $\rho_I$ | day<sup>-1</sup> | 0.1 | (3) | 0.1 | (3) | - |
| $\omega_{\rho_I}$ | day<sup>-1</sup> | 0.1 | (17) | 0.1 | (17) | - |
| $\mu_I$ | 10<sup>-5</sup> cell<sup>-1</sup> day<sup>-1</sup> | 2.9 | (18) | 2.9 | (18) | - |
| $\omega_{\mu_I}$ | 10<sup>-5</sup> cell<sup>-1</sup> day<sup>-1</sup> | 0.9 | (15) | 0.9 | (15) | - |
| *Residual errors* |  |  |  |  |  |  |
| $a_N$ | cell counts (log10) | 0.5 | (10) | 0.5 | (10) | - |
| $a_E$ | cell counts (log10) | 0.5 | (7) | 0.5 | (7) | - |
| $a_L$ | cell counts (log10) | 0.5 | (6) | 0.5 | (6) | - |
| $a_M$ | cell counts (log10) | 0.4 | (8) | 0.4 | (8) | - |

Figure 6 shows the estimated distribution for parameter *δ*_*NE*_ that varies within the population and for which we included a covariate. Histograms display the estimated individual parameter values of *δ*_*NE*_. They show two distinct distributions of *δ*_*NE*_ values, corresponding to VV (red)- and Tumor (green)-associated values. The histograms and the theoretical distributions are in agreement.

**Figure 6.**
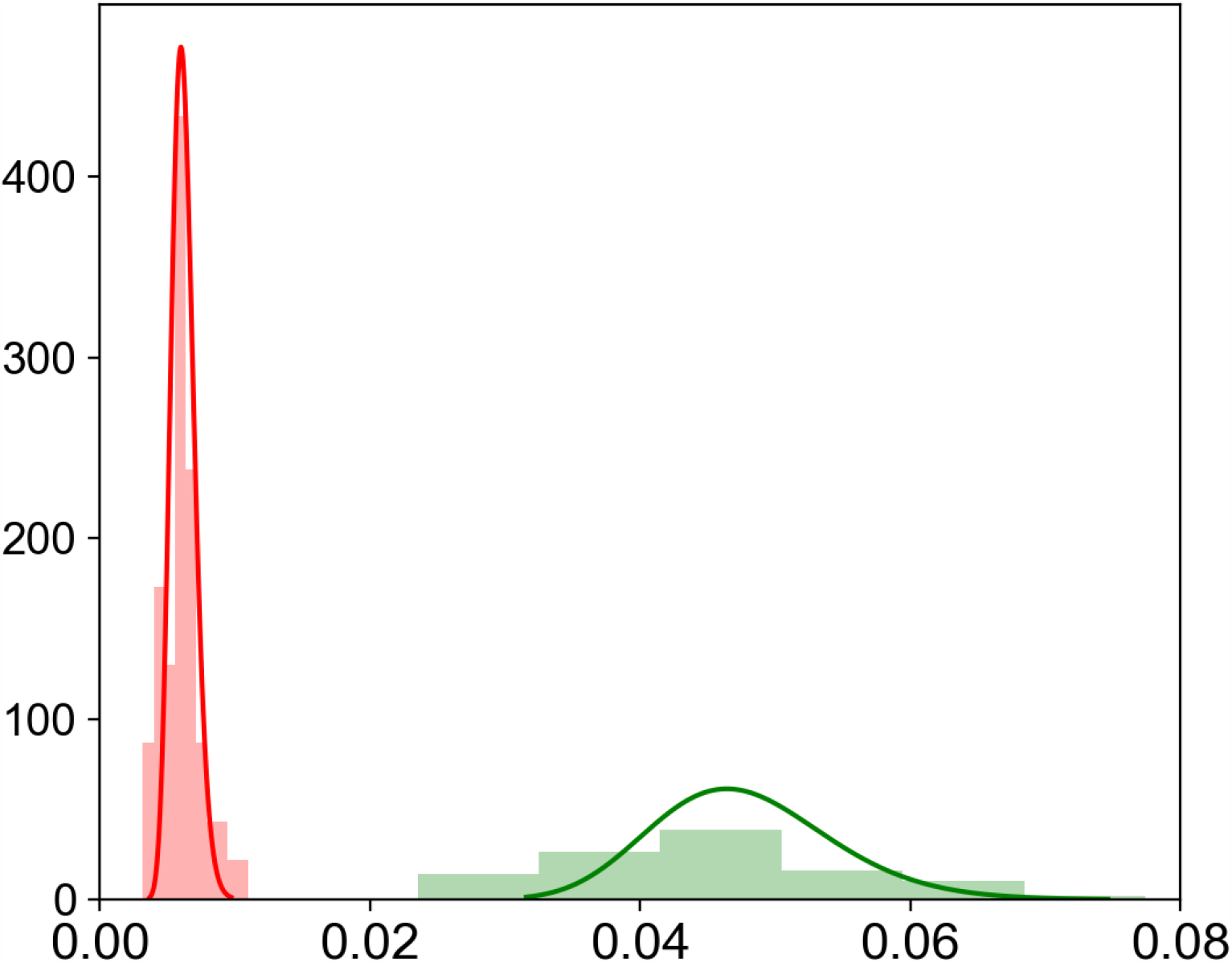
Probability distribution of parameter *δ*_*NE*_ defined with a covariate. Estimated distributions of VV-associated (left, red) and Tumor-associated (right, green) values are plotted. Histograms of estimated individual parameter values are also plotted (red for VV-associated values, green for Tumor-associated values).

Table 4 gives the estimated values of all parameters in both groups. Regarding parameters that do not vary within the population, it is required for parameters 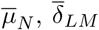 and 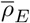 to be different to describe each data set, and this difference is accounted for with a covariate parameter. Noticeably, using categorical covariates mostly improves the confidence in the estimation, as highlighted by either RSE values in the same range 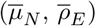 or improved (all other parameters) RSE values (Tables 3 and 4).

In summary, we identified parameters whose values are significantly different according to the immunogen used to activate CD8 T cells. These parameters correspond to the dynamics of naïve cells 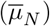, their activation (*δ*_*NE*_), the proliferation of early effector cells 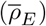, and differentiation to memory cells 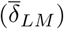. We hence conclude that different immunizations affect the CD8 T cell activation process in the first phase of the response (priming, activation of naive cells, expansion of the CD8 T cell population) as well as the development of the memory population. Different immunizations also induce various degrees of variability in the responses through the activation of naive cells, and our mathematical approach quantitatively estimates these degrees of variability.

### 2.4 Predicting dynamics following VV and Tumor immunizations

To challenge System (2) and the estimated parameters (Table 4), we compare simulated outputs to an additional data set, not used for data fitting up to this point, of both VV and Tumor immunizations, VV data set 2 and Tumor data set 2 (Table 1 and Section 4.8).

We already know the probability distribution of parameters (Table 4), so we only estimate individual parameters in order to fit individual dynamics. Results are shown in Figure 7, for both VV and Tumor data sets 2. Individual fits are available in Section 5.3. It is clear that estimated individual dynamics are consistent with previous individual dynamics estimations. Hence, we validate System (2) and values estimated in both VV and Tumor immunization contexts by showing that estimated parameter values allow to characterize CD8 T cell counts obtained in similar contexts as well as individual behaviors (Figure 7 and Section 5.3).

**Figure 7.**
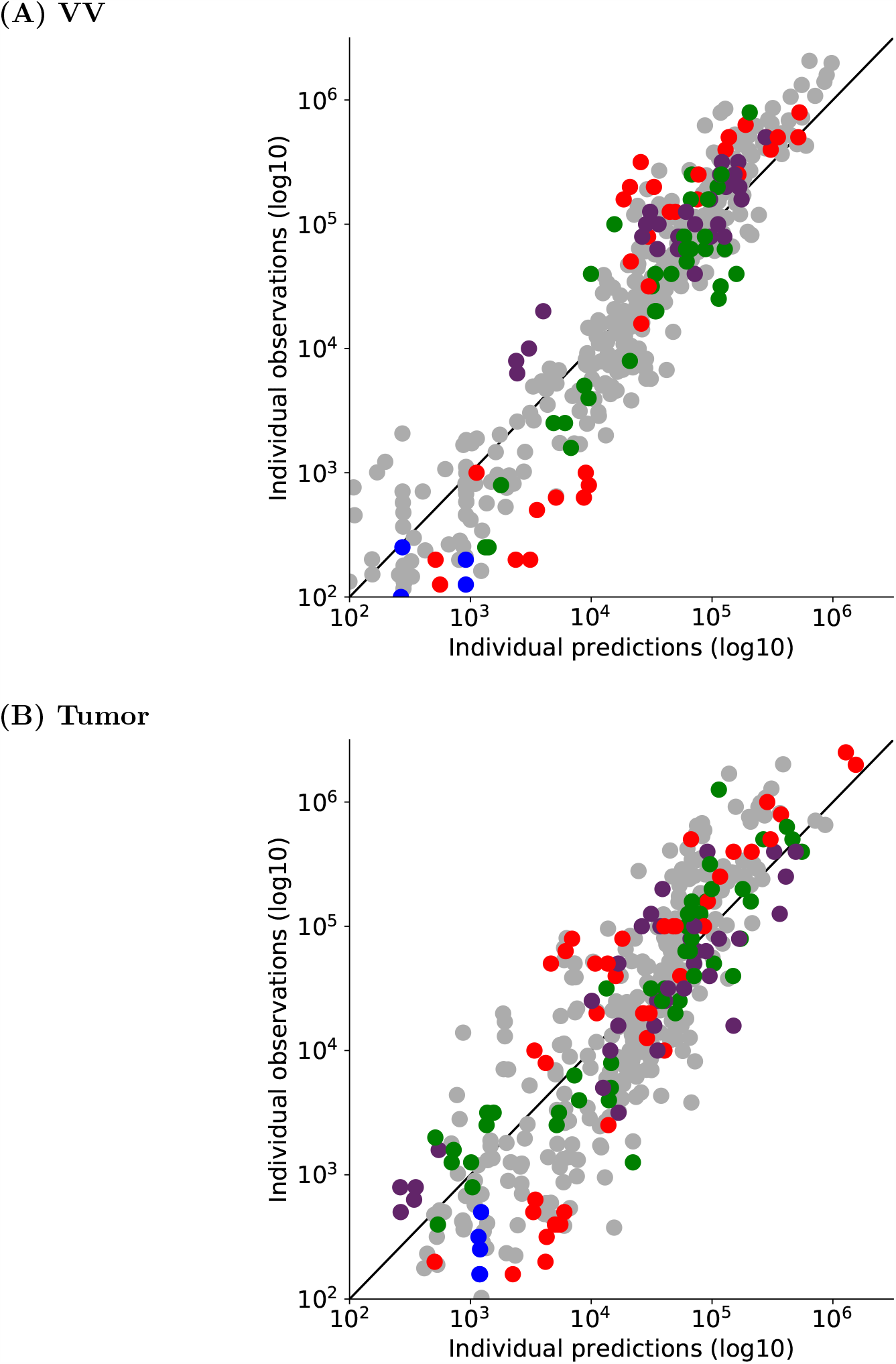
Observed *vs* estimated values of individual CD8 T cell counts for **(A)** VV data set 2 and **(B)** Tumor data set 2. Individual parameter values have been estimated with System (2) and population parameter values and distributions previously defined on VV and Tumor data set 1. In both figures, naive (blue), early effector (red), late effector (green), and memory (purple) cell counts are depicted. Grey points correspond to individual values from Figure 3.A and Figure 5.A. The black straight line is *y* = *x*.

## 3 Discussion

When following an *in vivo* immune response, experimental measurements are often limited by either ethical issues or tissue accessibility. Consequently, one often ends up measuring cell counts in peripheral blood on a restricted number of time points per individual, over the duration of a response (see Figure 4). Among measurements of a single individual, cell counts are often missing for one or more cell subpopulations. With such data, estimation of all model parameters becomes unlikely. Using nonlinear mixed effects models, we propose a dynamical model of CD8 T cell dynamics that circumvents this difficulty by assuming that all individuals within a population share the main characteristics. Using this framework, we propose an accurate description of individual dynamics, even though individual measurements are scarce. Indeed, we are able to obtain both good fits and relevant dynamics for individuals with only few cell count measurements, as illustrated in Figure 4. These results indicate that knowledge of population dynamics parameters and numerical simulations complement information given by experimental measurements.

Starting from the model described in Crauste et al. [10] that could efficiently describe CD8 T cell dynamics, at the level of average population cell-counts in peripheral blood, we built and validated this nonlinear mixed effects model in a step-wise fashion. The system was first modified to ensure correct parameter estimation when confronted to ideal, highly informative data. In a second step, the model was again modified and parameter values estimated by using experimental measurements generated through a VV immunization. We next identified parameters – hence biological processes – that vary between individuals and explain the between-individual variability, and other parameters that can be fixed within the population to explain biological data (measured in VV and Tumor immunization contexts). Finally, by including a categorical covariate we additionally identified immunization-dependent parameters.

In order to determine the contribution of each parameter to inter-individual variability, one could argue that performing a sensitivity analysis would shed light on the more sensitive parameters. Even though model’s output sensitivity to parameters and contribution of each parameter to individual variability may be partly related, they are nonetheless different concepts. Sensitivity analysis would highlight the influence of a parameter on the population dynamics, whereas our objective is to reproduce several individual outputs (individual dynamics) which exhibit more or less variability than the average population behavior. Therefore the use of classical tools like Sobol indices [31] or generalized sensitivity functions [32] is not adapted to handle the current question. Hence, we proposed a procedure, based on estimated errors and the shrinkage, to identify a minimal set of parameters (fixed and random effects) required to describe the data sets. It can be noticed that the shrinkage, expressed as a ratio of variances (see (5), provides an information similar to the one given by Sobol indices.

Noteworthy, from a biological point of view, the removal of one parameter during model reduction (for example, the death rate of late effector cells) must not be understood as if the corresponding process is not biologically meaningful. Rather, based on the available data, our methodology found that some processes are non-necessary in comparison with the ones described by the system’s equations.

Similarly, parameters characterizing immunogen dynamics vary within the population whereas model reduction led to remove the variability of equivalent processes (proliferation for instance) in CD8 T cell dynamics. It is likely that this is due to a lack of experimental measures on immunogen dynamics (whether virus load evolution or tumor growth), and one cannot conclude that inter-individual variability mostly comes from immunogen dynamics. Information on immunogen dynamics, when available, could significantly improve parameter estimation and help refining the information on inter-individual variability during CD8 T cell responses.

In our biological data, inter-individual variability is explained only by variability in the activation rate of naive cells, the mortality rate of effector cells, and dynamics (proliferation and death) of the immunogen. The former is actually in good agreement with the demonstration that in diverse infection conditions the magnitude of antigen-specific CD8 T cell responses is primarily controlled by clonal expansion [33].

Two of the three differentiation rates (early effector cell differentiation in late effector cells, and late effector cell differentiation in memory cells) do not need to vary to describe our data sets. This robustness of the differentiation rates is in good agreement with the auto-pilot model that shows that once naive CD8 T cells are activated their differentiation in memory cells is a steady process [34, 35].

Eventually, using nonlinear mixed effects models and an appropriate parameter estimation procedure, we were able to quantitatively reproduce inter-individual variability in two different immunization contexts (VV and Tumor) and provide predictive population dynamics when confronted to another data set (for both immunogens). This demonstrates the robustness of the model.

The addition of a categorical covariate allowed us to identify parameters that are immunization-dependent. Interestingly they control the activation of the response (priming, differentiation of naive cells, expansion of effector cells) as well as the generation of memory cells. This is again in good agreement with the biological differences that characterize the two immunogens used in this study. Indeed, pathogen-associated molecular patterns (PAMP) associated with vaccinia virus will activate a strong innate immune response that will provide costimulatory signals that in turn will increase the efficiency of naive CD8 T cell activation [5]. In contrast, when primed by tumor cells CD8 T cells will have access to limited amounts of costimulation derived from damage associated molecular patterns [36]. The amount of costimulation will also control the generation of memory cells [37]. Focusing on average CD8 T cell behaviors (not shown) highlights stronger responses following VV immunization, characterized by a faster differentiation of naive cells and a higher peak of the response (at approximately 3 × 10^5^ cells compare to 10^5^ cells for the Tumor induced response). Also, in average, more memory cells are produced following VV immunization. Hence the addition of covariates to the model parameters has allowed to identify biologically relevant, immunogen-dependent parameters.

Using covariates has additional advantages. First, they allow to consider a larger data set (in our case, the combination of two data sets) without adding too many parameters to estimate (4 covariates in our case). This is particularly adapted to situations where only some parameters are expected to differ depending on the data set (here, the immunogen). Second, and as a consequence, data fits may be improved compared to the situation where data sets generated with different immunogens are independently used to estimate parameters. Figure 8 illustrates this aspect: dynamics of two individuals are displayed, with and without covariate. In both cases using the covariate (and thus a larger data set) improved the quality of individual fits, and in the case of Individual 1 generated more relevant dynamics with a peak of the response occurring earlier, before day 10pi. No individual fit has been deteriorated by the use of a covariate (not shown).

**Figure 8.**
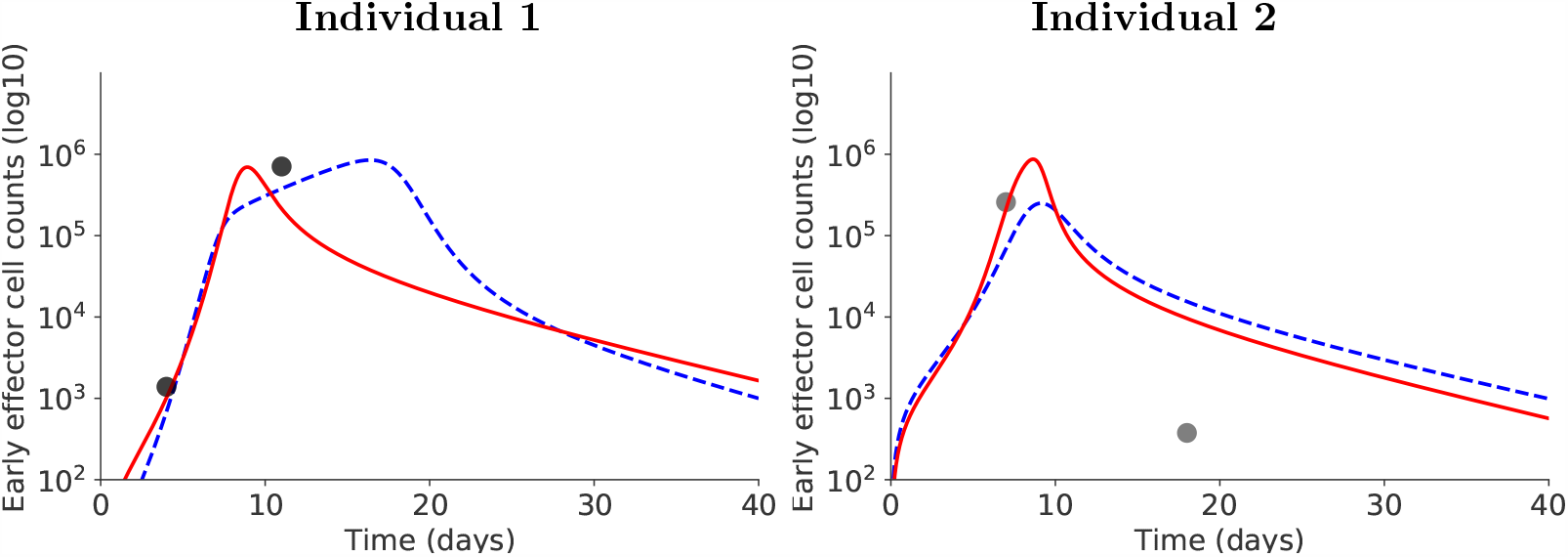
Positive side-effect of using covariates. For two illustrative individuals, accounting for covariates allows to better estimate early effector cell dynamics: red plain curve with covariate, blue dashed curve without covariate.

Finally, CD8 T cell response dynamics to both VV and Tumor immunogens were well captured for data sets that had not been used to perform parameter estimation (Section 2.4). The behavior of each individual was estimated with the prior knowledge acquired on the population (i.e. fixed parameters values and variable parameter distributions) and proved consistent with previous estimated individual behaviors. The correct prediction of individual behaviors by the model, in a simple mice experiment, paves the way to personalized medicine based on numerical simulations. Indeed, once the population parameters are defined, numerical simulation of individuals can be performed from a few measurements per individual and consequently would allow to adapt personalized therapies.

## 4 Material, Methods and Models

### 4.1 Ethics Statement

CECCAPP (Lyon, France) approved this research accredited by French Research Ministry under project #00565.01.

Mice were anesthetized either briefly by placement in a 3% isoflurane containing respiratory chamber or deeply by intraperitoneal injection of a mix of Ketamin (70 mg/kg) and Xylazin (9 mg/kg). All animals were culled by physical cervical disruption.

### 4.2 Data

All data used in this manuscript are available at https://osf.io/unkpt/?view_only=ff91bd89bc32421dbcbb356c3509ca55.

#### Experimental Models

C57BL/6 mice (C57BL6/J) and CD45.1+ C57BL/6 mice (B6.SJL-Ptprc^a^Pepc^b^/BoyCrl) were purchased from CRL. F5 TCR-tg mice recognizing the NP68 epitope were crossed to a CD45.1+ C57BL/6 background (B6.SJL-Ptprc^a^Pepc^b^/BoyCrl-Tg(CD2-TcraF5,CD2-TcrbF5)1Kio/Jmar) [38]. They have been crossed at least 13 times on the C57BL6/J background. All mice were homozygous adult 6-8-week-old at the beginning of experiments. They were healthy and housed in our institute’s animal facility under Specific Pathogen-Free conditions.

Age- and sex-matched litter mates or provider’s delivery groups, which were naive of any experimental manipulation, were randomly assigned to 4 experimental groups (of 5 mice each) and co-housed at least for one week prior to experimentation. Animals were maintained in ventilated enriched cages at constant temperature and hygrometry with 13hr/11hr light/dark cycles and constant access to 21 kGy-irradiated food and acid (pH = 3 ± 0.5) water.

#### Vaccinia Virus (VV) Immunization

2 × 10^5^ naive CD8 T cells from CD45.1+ F5 mice were transferred by retro-orbital injection in 59, 6-8-week-old, congenic CD45.2+ C57BL/6 mice briefly anaesthetized with 3% isoflurane. The day after deeply Xylazin/ Ketamin-anaesthetized recipient mice were inoculated intra-nasally with 2 × 10^5^ pfu of a vaccinia virus expressing the NP68 epitope (VV-NP68) provided by Pr. A.J. McMichael [38].

#### Tumor Immunization

2 × 10^5^ naive CD8 T cells from CD45.1+ F5 mice were transferred by retro-orbital injection in 55, 6-8-week-old, congenic CD45.2+ C57BL/6 mice briefly anaesthetized with 3% isoflurane. The day after, recipients were subcutaneously inoculated with 2.5 × 10^6^ EL4 lymphoma cells expressing the NP68 epitope (EL4-NP68) provided by Dr. T.N.M. Schumacher [39].

#### Phenotypic Analyses

Mice were bled at intervals of at least 7 days. Blood cell suspensions were cleared of erythrocytes by incubation in ACK lysis solution (TFS). Cells were then incubated with efluor780-coupled Fixable Viability Dye (eBioscience) to label dead cells. All surface stainings were then performed for 45 minutes at 4°C in PBS (TFS) supplemented with 1% FBS (BioWest) and 0.09% NaN3 (Sigma-Aldrich). Cells were fixed and permeabilized with the Foxp3-fixation and permeabilization kit (eBioscience) before intra-cellular staining for one hour to overnight. The following mAbs(clones) were utilized: Bcl2(BCL/10C4), CD45.1(A20) and CD45(30-F11) from Biolegend, Mki67(SolA15) and CD8(53.6.7) from eBioscience, and CD44 (IM7.8.1) from Miltenyi. Samples were acquired on a FACS LSR Fortessa (BD biosciences) and analyzed with FlowJo software (TreeStar).

#### CD8 T Cell Differentiation Stages

For both immunizations (VV and Tumor), phenotypic cell subsets based on Mki67-Bcl2 characterization [10] have been identified and the corresponding cell counts measured in blood, from day 4 post-inoculation (pi) up to day 47pi (VV and Tumor data sets 1, Table 1). Naive cells are defined as CD44-Mki67-Bcl2+ cells, early effector cells as CD44+Mki67+Bcl2-cells, late effector cells as CD44+Mki67-Bcl2-cells, and memory cells as CD44+Mki67-Bcl2+ cells [10].

### 4.3 Models of CD8 T cell dynamics

#### Initial model

The following system (3) is made of ODE and describes individual behaviors. This is the model in [10], it describes CD8 T cell subpopulation dynamics (see Section 4.2, Paragraph CD8 T Cell Differentiation Stages) as well as the immunogen load dynamics in primary immune responses, as follows

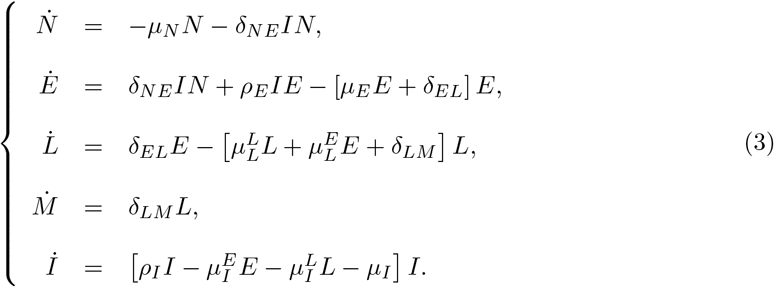

The variables *N, E, L* and *M* denote the four CD8 T cell subpopulation counts, naive, early effector, late effector, and memory cells respectively (see Section 4.2), and *I* is the immunogen load. For details regarding the construction and validation of this model we refer the readers to [10]. We hereafter briefly discuss this model.

The immunogen load dynamics are normalized with respect to the initial amount [10, 40], so *I*(0) = 1. The initial amounts of CD8 T cell counts are *N* (0) = 10^4^ cells, *E*(0) = 0, *L*(0) = 0 and *M* (0) = 0.

Parameters *δ*_*k*_ are the differentiation rates, with *k* = *NE, EL* or *LM* for differentiation from naive to early effector cells, from early effector to late effector cells and from late effector to memory cells, respectively.

Death parameters are denoted by *µ*_*k*_, where *k* = *N, E* and *I* for the death of naive cells, early effector cells and the immunogen respectively. Notations 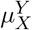 for some mortality-related parameters refer to parameters *µ*_*XY*_ in [10]: the subscript *X* refers to the CD8 T cell population or the immunogen that dies, and the superscript *Y* to the CD8 T cell population responsible for inducing death.

Early and late effector cells are cytotoxic, GrzB+ cells [10], so due to competition for limited resources (such as cytokines) and fratricidal death [41, 42] we assumed fratricide killing by CD8 T cells. Consequently the model accounts for effector-cell regulated death rates of both effector cells and the immunogen [10, 40]. Natural mortality rates are considered for naive and memory cells (*µ*_*N*_, *µ*_*I*_).

Proliferation parameters of early effector cells and the immunogen are respectively denoted by *ρ*_*E*_ and *ρ*_*I*_. Proliferation of both CD8 T cells and the immunogen are partially controlled by the immunogen, so proliferation rates are assumed to depend on *I*. Noticeably, among CD8 T cells only early effector cells are Mki67+ cells so they are the only cells assumed to proliferate and divide [10].

System (3) has been introduced and validated on a similar VV data set in [10]. To account for individual behavior, parameters will be complexified assuming they are drawn from probability distributions and in the same time this system will be simplified through a model selection procedure to ensure the correct estimation of parameter values with available data sets (see Sections 4.6 and 4.7).

#### Model selected on synthetic data

Model (3) has been obtained by fitting average dynamics of a CD8 T cell immune response [10]. When confronting this model to heterogeneous data of individual CD8 T cell dynamics and using mixed effects modeling, it is mandatory to verify that assumptions of the mixed effects model (see Section 4.4) are valid. To investigate this mathematical property, we will rely on synthetic data that are highly informative compared to experimental data and, additionally, for which we know the parameter values behind data generation so we possess an explicit control on parameter estimations. Using synthetic data and the procedure described in Section 4.6 leads to the selection of the System (1),

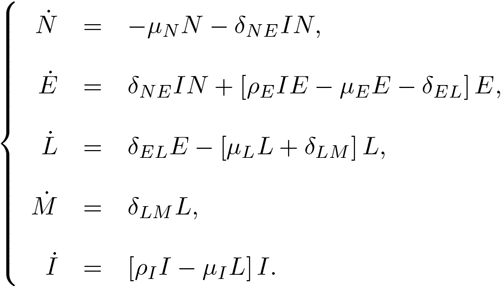

This model is dynamically similar to System (3), but in order to correctly fit synthetic data and to satisfy the assumptions of mixed effects modeling, parameters 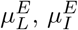 and *µ*_*I*_ have been removed: it was not possible to accurately estimate them to non-zero true values. For the sake of simplicity the parameters are renamed in System (1): 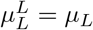 and 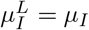 System (1) is defined by 9 parameters.

#### Model selected on biological data

When using biological, *in vivo* experimental data instead of synthetic data, not as many measurements per individual can be obtained (see Table 1) so the dynamical model may easily be over-informed (too many parameters compared to the size of the sampling). Using System (1), the confrontation with VV data set 1 leads to the modified System (2),

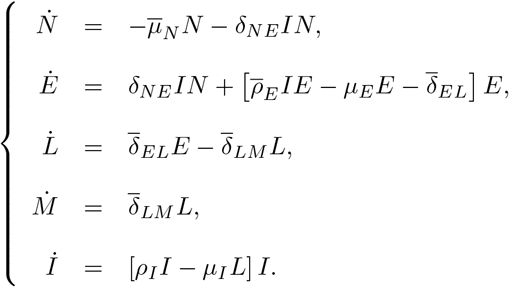

System (2) has 8 parameters (*µ*_*L*_ has been removed from System (1)): 4 parameters are fixed within the population 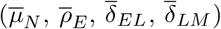 and 4 parameters have a random effect (*δ*_*NE*_, *µ*_*E*_, *ρ*_*I*_, 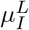).

### 4.4 Nonlinear mixed effects models

Nonlinear mixed effects models allow a description of inter-individual heterogeneity within a population of individuals (here, mice). The main idea of the method is to consider that since all individuals belong to the same population they share common characteristics. These common characteristics are called “fixed effects” and characterize an average behavior of the population. However, each individual is unique and thus differs from the average behavior by a specific value called “random effect”.

This section briefly describes our main hypotheses. Details on the method can be found in [17–19, 43].

Each data set {*y*_*i,j*_, *i* = 1, …, *N*_*ind*_, *j* = 1, …, *n*_*i*_} is assumed to satisfy

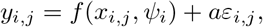

where *y*_*i,j*_ is the *j*^*th*^ observation of individual *i, N*_*ind*_ is the number of individuals within the population and *n*_*i*_ is the number of observations for the *i*^*th*^ individual.

The function *f* accounts for individual dynamics generated by a mathematical model. In this work *f* is associated with the solution of a system of ODE, see Section 4.3. The function *f* depends on known variables, denoted by *x*_*i,j*_, and parameters of the *i*^*th*^ individual, denoted by *ψ*_*i*_.

Individual parameters *ψ*_*i*_ are assumed to be split into fixed effects (population-dependent effects, average behavior) and random effects (individual-dependent effects). If 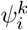 denotes the *k*-th parameter characterizing individual *i*, then it is assumed that

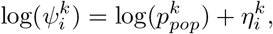

where the vector of parameters 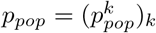 models the average behavior of the population, and 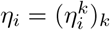 represents how the individual *i* differs from this average behavior. Variables 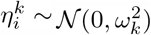, and they are assumed independent and identically distributed. The variance 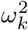 quantifies the variability of the *k*-th parameter within the population. From now on we will denote by *ω*^2^ the vector of variances 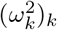. Parameters *ψ*_*i*_ are assumed to follow a log-normal distribution to ensure their positivity.

The residual errors, combining model approximations and measurement noise, are denoted by *aε*_*i,j*_. They quantify how the model prediction is close to the observation. Residual errors are assumed independent, identically and normally distributed, *i*.*e ε*_*i,j*_ ∼ 𝒩 (0, 1). Moreover, the random effects *η*_*i*_ and the residual errors *aε*_*i,j*_ are mutually independent. In this work, we assume a *constant* error model, with a constant *a*, for all cell populations, since they are all observed in log10 scale. The error parameter is estimated for each subpopulation (naive cells - *a*_*N*_; early effector cells - *a*_*E*_; late effector cells - *a*_*L*_; memory cells - *a*_*M*_). When data on the immunogen dynamics are available (only when using synthetic data), we assume a proportional error for the immunogen which is observed, so that *a*_*I*_ = *b*_*I*_*f*.

We will write that a parameter is *fixed within the population* if all individuals have the same value for this parameter. On the contrary, if the variance 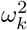 of a parameter is non-zero, then this parameter will account for inter-individual variability within the population.

### 4.5 Parameter estimation

Parameter values are estimated with the Stochastic Approximation Expectation-Maximization (SAEM) algorithm. This algorithm is adapted to non-linear mixed effects models [18] and has been shown to quickly converge under general conditions [17]. Moreover, an implementation of the SAEM algorithm is available in Monolix [30], a freely available software that provides different indicators to quantify the quality of estimations and fit. We used the SAEM algorithm and Monolix in this work.

#### Population and individual parameters

Under the previous assumptions (Section 4.4), cell population dynamics (average behavior and inter-individual variability) are described by parameters: *p*_*pop*_, *ω*^2^ and *a*. These parameters are estimated by maximizing the likelihood with the SAEM algorithm.

Once these parameters have been estimated, each individual vector of parameters *ψ*_*i*_ is estimated by maximizing the conditional probabilities 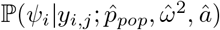, where 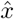 denotes the estimated value of *x*.

Both estimations are performed with Monolix software [30]. Files to run the algorithm (including all algorithm parameters) are available at https://plmlab.math.cnrs.fr/audebert/cd8-responses.

#### Covariates

In order to study whether differences observed in parameter values between VV and Tumor data sets (Table 1) are only related to random sampling or if they can be explained by the immunogen, we use categorical covariates (Section 2.3).

To tackle this question, we first pool together VV and Tumor data sets 1. Second, using this full data set, we estimate parameter values by assuming that fixed effects of some Tumor-associated parameters are different from those of the corresponding VV-associated parameters.

To introduce categorical covariates in our mixed effect model, we assume that if an individual is either in Tumor or VV data set then the probability distribution of its individual parameter vector *ψ*_*i*_ has a different mean. We write

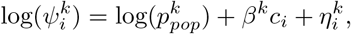

where *c*_*i*_ equals 0 if individual *i* is in VV data set 1 and 1 if it is in Tumor data set 1, and *β* = (*β*^*k*^)_*k*_ is a vector of covariate parameters. We test whether the estimated covariate parameter 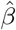 is significantly different from zero with a Wald test, using [30] software, and we use a *p*-value threshold at 0.05.

Parameters (*p*_*pop*_, *ω*^2^, *a, β*) are then characterizing cell population dynamics for both VV and Tumor immunogens. If the estimated vector 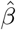 is significantly different from zero, then part of the experimentally observed variability could be explained by the immunogen.

### 4.6 Model selection on synthetic data

Model selection relies on criteria that allow to evaluate to which end a model appropriately satisfies *a priori* assumptions. For instance, one usually requires a model to correctly fit the data, and uses so-called quality of fit criteria, and/or requires that initial modeling assumptions are satisfied.

Here, we do not use quality of fit criteria to select a model because all models correctly fit data due to a priori over-informed models that have too many parameters compared to available data (see Paragraph Model selection below). Instead, we focus on the capacity of the parameter estimation procedure to correctly estimate model parameters and to the a posteriori validation of statistical assumptions. To do so, we first use synthetic data (see Paragraph Generation of synthetic data below). We take advantage of the fact that we know the exact parameter values used to generate synthetic data, so in order to evaluate the correctness of estimated parameter values we rely on:

- the relative difference between the estimated parameter value and the true value,
- the relative standard error (RSE), defined as the ratio between the standard error (square root of the diagonal elements of the variance-covariance matrix) and the estimated value of the parameter [19],

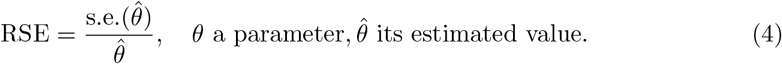 A large RSE indicates a poor estimation of the parameter.
- the *η*-shrinkage value (denoted throughout this manuscript as the *shrinkage* value), defined as

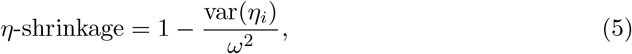

where var(*η*_*i*_) is the empirical variance of the random effect *η*_*i*_ and *ω*^2^ the estimated variance of the parameter; Large values of the shrinkage characterize individual estimates shrunk towards the conditional mode of the parameter distribution.

We decided not to consider the mathematical notion of identifiability here. Indeed, studying identifiability in nonlinear mixed effect models is a complicated, open question that has been discussed for instance in [44]. Approaches based on the Fisher Information Matrix (RSE) have been proposed and are often used for evaluating identifiability of population parameters, while analysis of the shrinkage allows to investigate individual parameters identifiability, and we used such methods in this work.

#### Generation of synthetic data

Using a dynamical model (here System (3)), we generate a set of data associated to solutions of the model, where all the parameters are drawn from known log-normal distributions. Parameters *p*_*k*_ varying in the population satisfy log(*p*_*k*_) ∼ 𝒩 (log(*m*_*k*_), 0.1^2^). The standard deviation is fixed to the value 0.1 to generate heterogeneity, and values of medians *m*_*k*_ are given in Table A.1. A multiplicative white noise modifies model’s outputs in order to mimic real measurements (we consider a white noise with standard deviation 0.2).

These data consist of time points and measurements for the 4 subpopulations of CD8 T cell counts (in log10 scale) and the immunogen load. These are called *synthetic data*, and these sets of data are referred to as Synth data set X, with X= 1, …, 4 (Table 1).

We generate synthetic data for 100 individuals, cell counts are sampled at days 4, 5, 6, 7, 8, 9, 10, 12, 14, 16, 18, 20, 25, 30pi (cf. Figure A.1 to A.4). In agreement with real biological data, we assume that all cell counts below 100 cells are not measured, and remove the data. For the immunogen load, values lower than 0.1 are also not considered.

#### Model selection

Model selection on synthetic data is performed in 4 steps:

**Step 1** Select an initial model

**Step 2** Estimate parameter values using SAEM [30]

**Step 3** Remove (priority list):

- parameters whose estimated value is different from their true value, and the RSE is larger than 5%
- random effect of parameters with shrinkage larger than 30%.

**Step 4** Select a model with all parameters correctly estimated

In Step 1, model (3) is used, with all parameters varying within the population. This makes 29 parameters to estimate: 12 mean values, 12 random effects, 5 error parameters.

In Step 3, based on the estimations performed in Step 2, we iteratively remove parameters that are not correctly estimated. To do so, we first focus on parameters that are not estimated to their true value (which is known) and whose RSE is larger than 5% (this threshold corresponds to a 5% error on the estimated value, see (4). We consider that the estimated value is different from the true value if *ε*_*rr*_ *>* 10%, with

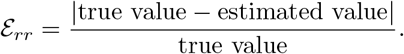

Once all parameters are correctly estimated according to the two first criteria, we remove random effects of parameters with shrinkage larger than 30% (Savic and Karlsson [45] have shown that shrinkage can generate false correlations between random effects, or mask the existing correlations, starting from 30% shrinkage).

One must note that every time a parameter is removed from the model (mean value or random effect) then new synthetic data are generated using the same protocol as described above, and Step 2 is performed again.

Errors are known when using synthetic data: since a normal noise, proportional to the observation, modifies each observation then there is a constant error on observations of cell counts in log10 scale, and a proportional error on the immunogen load. As mentioned in Section 4.4, we assume a constant error for all cell populations and a proportional error for the immunogen load. Diagnostic tools in [30] show that error models are correct (not shown here).

Quality of fit criteria do not provide relevant information in our case: the Bayesian Information Criterion (BIC) reaches very low values, even for the initial model (3), whereas observations *vs* predictions graphs show that the number of outliers is not modified by simplifications of the model. Hence, we do not use quality of fit criteria to select a model. In Step 4, we select a model based on the chosen criteria that insures the correct estimation of all its parameters and its reduced shrinkage when confronted to a set of synthetic data.

### 4.7 Model selection on biological data

Biological data are the ones introduced in Section 4.2. Compared to synthetic data, we do not know the parameter values that would characterize them and they provide less observations, hence it may not be possible to correctly estimate as many parameters as in the synthetic data case.

Model selection on biological data is also performed in 4 steps:

**Step 1** Select an initial model

**Step 2** Estimate parameter values using SAEM [30]

**Step 3** Remove (priority list):

- parameters whose RSE is larger than 100%
- random effect of parameters with shrinkage larger than 75%

**Step 4** Select a model with RSE and shrinkages low

In Step 1, model (1) is used, with all parameters varying within the population. This makes 23 parameters to estimate: 9 mean values, 9 random effects, 5 error parameters. This model is the one selected on synthetic data (see Section 2.1).

In Step 3, we iteratively remove parameters that are not correctly estimated. We first focus on parameters that are not estimated with a high confidence, that is RSE *>* 100%. Once all parameters are correctly estimated, we remove random effects of parameters with shrinkage larger than 75%. Noticeably, we cannot use the same threshold values for the RSE and the shrinkage when using either synthetic or real data, because measurement errors are different: controlled and known for synthetic data, uncontrolled and a priori unknown for real data, with measurement uncertainties.

The error model is not known, so we use the same error model as for synthetic data: a constant error for all cell populations (note that no data on immunogen is available, so the error parameter for the immunogen is not estimated). Diagnostic tools in [30] show that assuming constant error models is acceptable (not shown here).

### 4.8 A posteriori model validation on biological data

In Section 2.4, the model selected on biological data is compared to data that were not used for parameter estimation. These data are presented hereafter.

In order to assess the model ability to characterize and predict immune response dynamics we compare our results to additional experiments, VV data set 2 and Tumor data set 2 (see Table 1 and Section 4.2), similar to the ones used to estimate parameters (VV and Tumor data sets 1). CD8 T cell counts of naive, early and late effector, and memory cells have been measured following VV and Tumor immunizations, on days 4, 6, 7, 8, 11, 13, 15, 21, 28, 42pi.

The probability distribution of parameters (mean values, random effects) are known since we have estimated them on VV and Tumor data sets 1 (Section 4.7). These parameters are not estimated on the validation data. We use them to estimate the individual parameter values that fit individual behaviors of these new data sets (see Section 4.5).

## 5 Appendices

### 5.1 Parameter values used to generate synthetic data sets

Table A.1 lists parameter values used to generate Synth data sets 1 to 4 (see Table 1). Figures A.1 to A.4 illustrate Synth data sets 1 to 4 kinetics.

#### 5.2 Parameter value estimation using Synth data sets 1 to 4

Table A.2 presents the different steps in estimating parameter values using Synth data sets 1 to 4 and System (3). The procedure is detailed in Section 4.6.

#### 5.3 Predicted individual dynamics from VV and Tumor data sets 2

Predicted individual dynamics from VV and Tumor data sets 2, discussed in Section 2.4, are available at https://plmlab.math.cnrs.fr/audebert/cd8-responses.

## Acknowledgments

The authors are grateful to Pr. Adeline Leclercq Samson for sharing her expertise on nonlinear mixed effects models. We thank the BioSyL Federation and the LabEx Ecofect (ANR-11-LABX-0048) of the University of Lyon for inspiring scientific events. This work was supported by Inria PRE MEMOIRE grant and by the ANR predivac grant (ANR-12-RPIB-0011-01). We acknowledge the contributions of SFR Biosciences (UMS3444/CNRS, US8/Inserm, ENS de Lyon, UCBL) and of the CELPHEDIA Infrastructure (http://www.celphedia.eu/), especially the center AniRA in Lyon (AniRA-PBES and AniRA-Cytometrie facilities).

## Author Contributions

Chloe Audebert designed the model, performed all simulations and analyses, and wrote the manuscript. Daphne Laubreton performed the experimental work, reviewed and edited the manuscript. Christophe Arpin conceived the study and designed the experiments, reviewed and edited the paper. Olivier Gandrillon contributed to models’ analyses, reviewed and edited the manuscript. Jacqueline Marvel conceived the study, designed the experiments, reviewed and edited the manuscript. Fabien Crauste helped designing the model and analyzing the results, wrote the paper, and secured the funding.

---

**Table A1.** Parameter values of fixed effects (median values) used to generate Synth data sets 1 to 4 from System (3) and its subsequent reductions: removal of 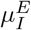 (column 4), of 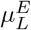 (column 5), and of *µ*_*I*_ (column 6). Notations 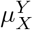 for some mortality-related parameters refer to parameters *µ*_*XY*_ in [10]: the subscript *X* refers to the CD8 T cell population or the immunogen that dies, and the superscript *Y* to the CD8 T cell population responsible for inducing death.

| Parameter | Unit | Synth data set |  |  |  |
| --- | --- | --- | --- | --- | --- |
|  |  | 1 | 2 | 3 | 4 |
| $\mu_N$ | $\text{day}^{-1}$ | 0.75 | 0.75 | 0.75 | 0.75 |
| $\delta_{NE}$ | $\text{day}^{-1}$ | 0.009 | 0.009 | 0.009 | 0.009 |
| $\rho_E$ | $\text{day}^{-1}$ | 0.64 | 0.64 | 0.64 | 0.64 |
| $\mu_E^E$ | $10^{-6} \text{ cell}^{-1} \text{ day}^{-1}$ | 21.5 | 21.5 | 21.5 | 21.5 |
| $\delta_{EL}$ | $\text{day}^{-1}$ | 0.59 | 0.59 | 0.59 | 0.59 |
| $\mu_L^L$ | $10^{-6} \text{ cell}^{-1} \text{ day}^{-1}$ | 7.5 | 7.5 | 7.5 | 7.5 |
| $\mu_L^E$ | $10^{-8} \text{ cell}^{-1} \text{ day}^{-1}$ | 3.6 | 3.6 | - | - |
| $\delta_{LM}$ | $\text{day}^{-1}$ | 0.025 | 0.025 | 0.025 | 0.025 |
| $\rho_I$ | $\text{day}^{-1}$ | 0.1 | 0.1 | 0.1 | 0.1 |
| $\mu_I^E$ | $10^{-7} \text{ cell}^{-1} \text{ day}^{-1}$ | 1.8 | - | - | - |
| $\mu_I^L$ | $10^{-5} \text{ cell}^{-1} \text{ day}^{-1}$ | 1.8 | 1.8 | 1.8 | 1.8 |
| $\mu_I$ | $\text{day}^{-1}$ | 0.055 | 0.055 | 0.055 | - |

**Table A2.** Steps in estimating parameter values using Synth data sets 1 to 4 and System (3). The procedure is detailed in Section 4.6. True values of parameters (fixed effects) are given on the second line, true values of random effects all equal 0.1. At *Step 1*, the procedure leads to removing parameter 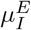. At *Step 2*, the procedure leads to removing parameter 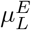. At *Step 3*, the procedure leads to removing parameter *µ*_*I*_. At *Step 4*, no other action is required. Values used to take a decision are highlighted in bold at each step. In the first column, ‘m.v.’ stands for mean value, RSE is defined in (4), ‘r.e.’ stands for random effect, and the shrinkage is defined in (5). Note that values (mean values and random effects) of parameters 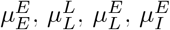 and 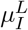 have to be multiplied by 10^*−*5^ (for 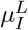), 10^*−*6^ (for 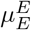 and 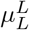), 10^*−*7^ (for 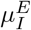), and 10^*−*8^ (for 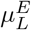). Units are omitted for the sake of clarity.

| Parameters | $\mu_N$ | $\delta_{NE}$ | $\rho_E$ | $\mu_E^E$ | $\delta_{EL}$ | $\mu_L^L$ | $\mu_L^E$ | $\delta_{LM}$ | $\rho_I$ | $\mu_I^E$ | $\mu_I^L$ | $\mu_I$ |
| --- | --- | --- | --- | --- | --- | --- | --- | --- | --- | --- | --- | --- |
| True values | 0.75 | 0.009 | 0.64 | 21.5 | 0.59 | 7.5 | 3.6 | 0.025 | 0.1 | 1.8 | 1.8 | 0.055 |
| *Step 1* |  |  |  |  |  |  |  |  |  |  |  |  |
| m.v. | 0.75 | 0.009 | 0.60 | 15.8 | 0.59 | 7.5 | 0.9 | 0.024 | 0.07 | **0.2** | 1.8 | 0.012 |
| RSE | 1 | 2 | 2 | 4 | 2 | 2 | 21 | 3 | 2 | **61** | 3 | 11 |
| r.e. | 0.1 | 0.2 | 0.2 | 0.3 | 0.2 | 0.2 | 0.7 | 0.3 | 0.2 | 2 | 0.3 | 0.9 |
| RSE | 7 | 7 | 7 | 8 | 7 | 7 | 23 | 7 | 8 | 21 | 7 | 11 |
| shrinkage | -1 | 8 | 12 | 18 | 13 | 1 | 100 | -1 | 39 | 96 | 4 | 82 |
| *Step 2* |  |  |  |  |  |  |  |  |  |  |  |  |
| m.v. | 0.76 | 0.009 | 0.59 | 17.2 | 0.58 | 7.9 | **0.3** | 0.024 | 0.07 | - | 1.9 | 0.012 |
| RSE | 1 | 2 | 2 | 4 | 2 | 2 | **17** | 3 | 2 | - | 3 | 11 |
| r.e. | 0.1 | 0.2 | 0.2 | 0.3 | 0.2 | 0.2 | 0.5 | 0.3 | 0.2 | - | 0.3 | 0.9 |
| RSE | 7 | 7 | 7 | 8 | 7 | 7 | 26 | 7 | 8 | - | 7 | 10 |
| shrinkage | -1 | 10 | 12 | 20 | 15 | 1 | 100 | -1 | 46 | - | 4 | 80 |
| *Step 3* |  |  |  |  |  |  |  |  |  |  |  |  |
| m.v. | 0.75 | 0.01 | 0.59 | 17.0 | 0.58 | 7.8 | - | 0.025 | 0.07 | - | 1.8 | **0.013** |
| RSE | 1 | 2 | 2 | 3 | 2 | 2 | - | 3 | 3 | - | 3 | **9** |
| r.e. | 0.1 | 0.2 | 0.2 | 0.3 | 0.2 | 0.2 | - | 0.3 | 0.2 | - | 0.3 | 0.7 |
| RSE | 7 | 7 | 7 | 8 | 7 | 7 | - | 7 | 8 | - | 7 | 10 |
| shrinkage | -1 | 8 | 11 | 17 | 13 | 1 | - | -1 | 34 | - | 2 | 82 |
| *Step 4* |  |  |  |  |  |  |  |  |  |  |  |  |
| m.v. | 0.76 | 0.009 | 0.66 | 21.2 | 0.61 | 7.3 | - | 0.024 | 0.1 | - | 1.8 | - |
| RSE | 1 | 2 | 1 | 2 | 1 | 2 | - | 3 | 1 | - | 2 | - |
| r.e. | 0.1 | 0.2 | 0.1 | 0.2 | 0.1 | 0.2 | - | 0.3 | 0.1 | - | 0.2 | - |
| RSE | 7 | 7 | 7 | 7 | 7 | 7 | - | 7 | 7 | - | 7 | - |
| shrinkage | -1 | 3 | 6 | 6 | 7 | -1 | - | -1 | 0 | - | 1 | - |

**Figure A1.**
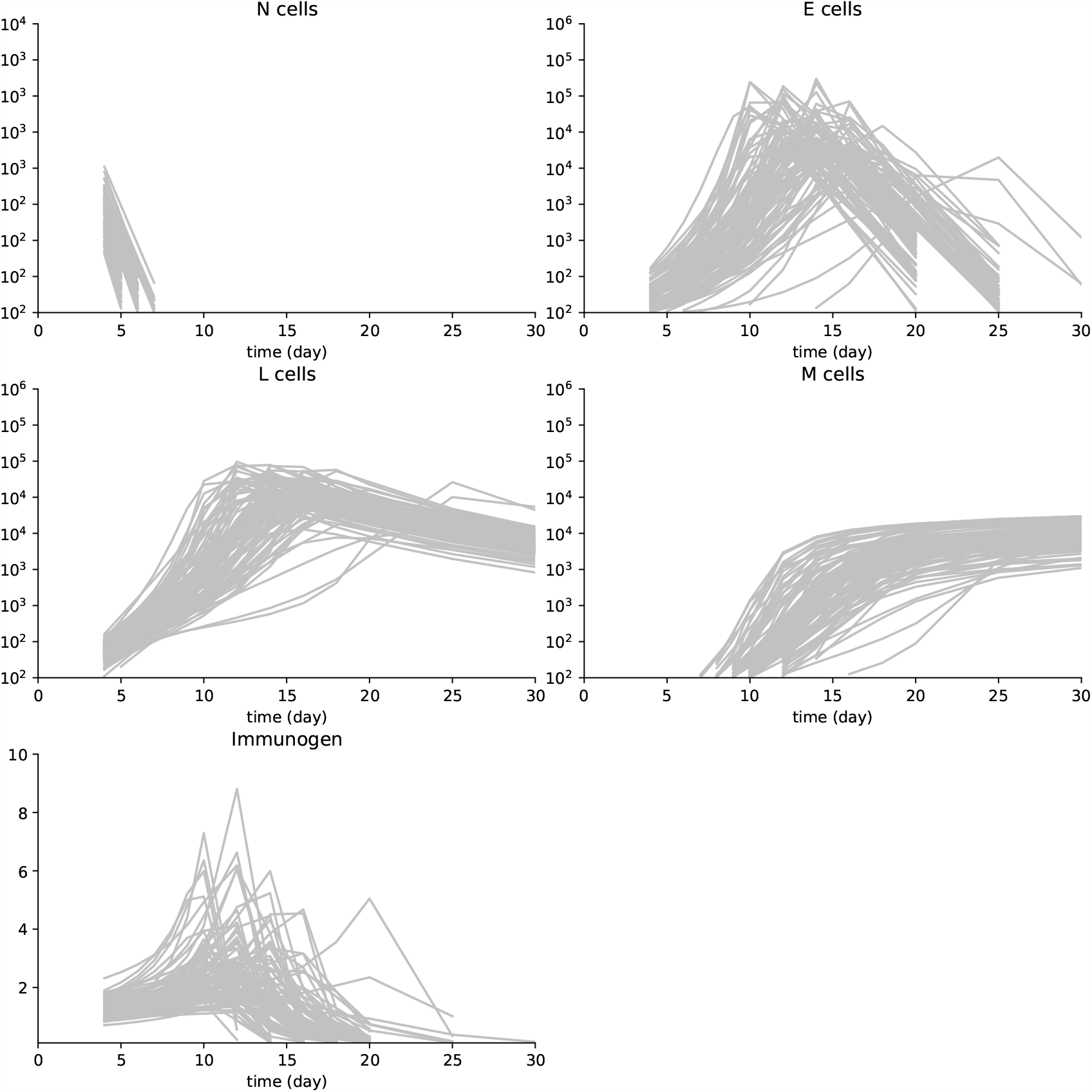
Synth data set 1. These data have been obtained by simulating System (3) with parameter values in Table A.1 and using a multiplicative white noise, as detailed in Section 4.6. 100 individuals are simulated and first observations are on day 4 pi for cell populations and the immunogen. Then measurements are on days 5, 6, 7, 8, 9, 10, 12, 14, 16, 18, 20, 25, and 30 pi. All cell counts below 100 cells are not measured. For the immunogen load, values lower than 0.1 are also not considered.

**Figure A2.**
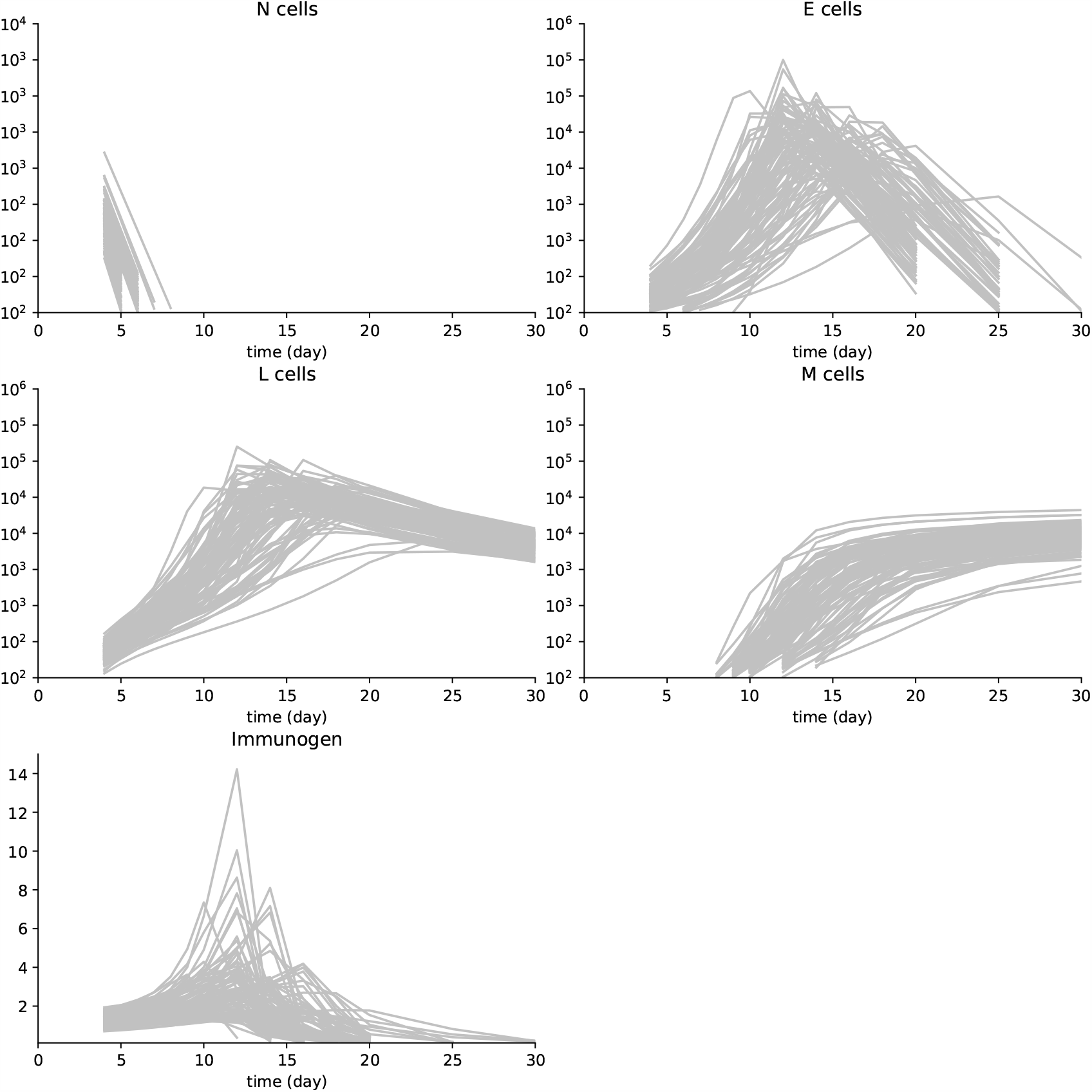
Synth data set 2. These data have been obtained by simulating a reduced System (3), with parameter values in Table A.1, and using a multiplicative white noise, as detailed in Section 4.6. See Figure A.1 for details.

**Figure A3.**
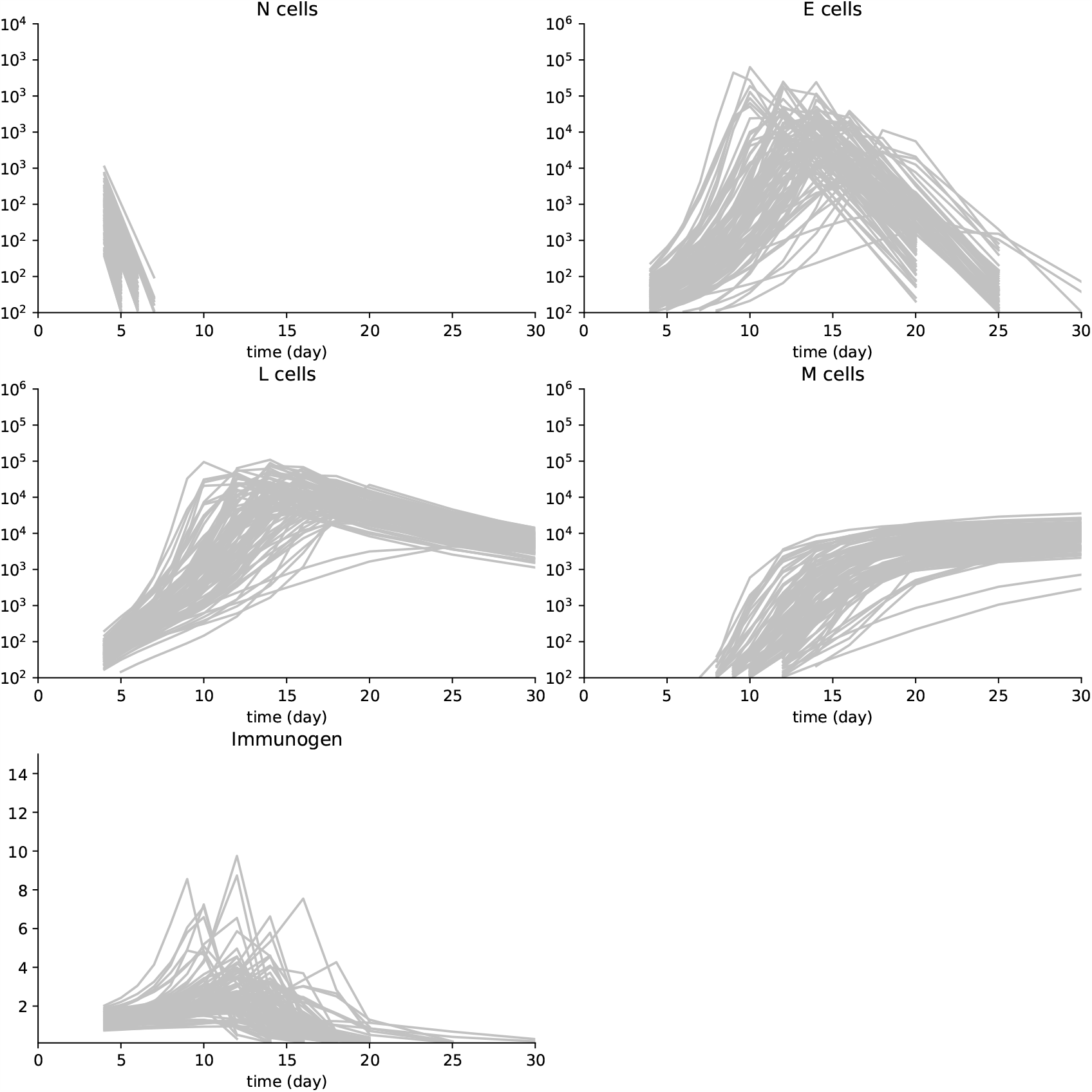
Synth data set 3. These data have been obtained by simulating a reduced System (3), with parameter values in Table A.1, and using a multiplicative white noise, as detailed in Section 4.6. See Figure A.1 for details.

**Figure A4.**
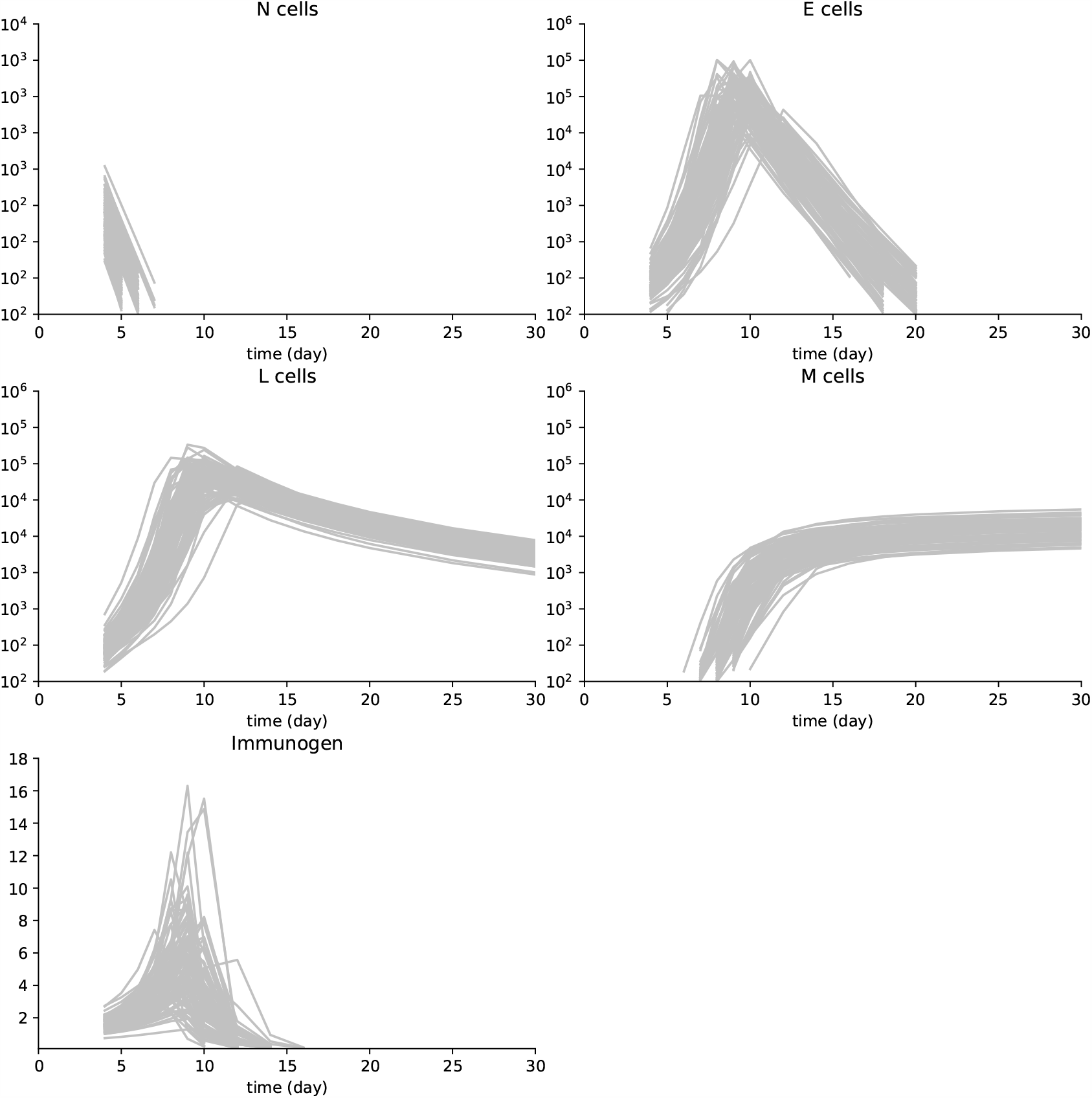
Synth data set 4. These data have been obtained by simulating a reduced System (3), with parameter values in Table A.1, and using a multiplicative white noise, as detailed in Section 4.6. See Figure A.1 for details.

## Notes

https://osf.io/unkpt/?view_only=ff91bd89bc32421dbcbb356c3509ca55

https://plmlab.math.cnrs.fr/audebert/cd8-responses

### Competing Interest Statement

The authors have declared no competing interest.

### Summary of Updates

We updated the description of our model selection strategy and the parameter estimation procedures. All our main results remain valid.

